# Viroid-like colonists of human microbiomes

**DOI:** 10.1101/2024.01.20.576352

**Authors:** Ivan N. Zheludev, Robert C. Edgar, Maria Jose Lopez-Galiano, Marcos de la Peña, Artem Babaian, Ami S. Bhatt, Andrew Z. Fire

## Abstract

Here, we describe the “Obelisks,” a previously unrecognised class of viroid-like elements that we first identified in human gut metatranscriptomic data. “Obelisks” share several properties: (i) apparently circular RNA ∼1kb genome assemblies, (ii) predicted rod-like secondary structures encompassing the entire genome, and (iii) open reading frames coding for a novel protein superfamily, which we call the “Oblins”. We find that Obelisks form their own distinct phylogenetic group with no detectable sequence or structural similarity to known biological agents. Further, Obelisks are prevalent in tested human microbiome metatranscriptomes with representatives detected in ∼7% of analysed stool metatranscriptomes (29/440) and in ∼50% of analysed oral metatranscriptomes (17/32). Obelisk compositions appear to differ between the anatomic sites and are capable of persisting in individuals, with continued presence over >300 days observed in one case. Large scale searches identified 29,959 Obelisks (clustered at 90% nucleotide identity), with examples from all seven continents and in diverse ecological niches. From this search, a subset of Obelisks are identified to code for Obelisk-specific variants of the hammerhead type-III self-cleaving ribozyme. Lastly, we identified one case of a bacterial species (*Streptococcus sanguinis*) in which a subset of defined laboratory strains harboured a specific Obelisk RNA population. As such, Obelisks comprise a class of diverse RNAs that have colonised, and gone unnoticed in, human, and global microbiomes.

## Introduction

RNA viruses (*Riboviria*) are in part defined by their encoding of their own replicative polymerases, a feature that can be leveraged for homology-based viral discovery ^1–5^. By contrast, viroids ^6,7^ and Hepatitis Delta-like viral (HDV) ‘satellites’ ^8^ (Supplementary Figure 1) co-opt eukaryotic host RNA polymerases for their replication, resulting in some of biology’s smallest known genomes (viroids: ∼350 nt; Delta: ∼1.7 kb). These streamlined genomes define the working limits of biological information transfer ^9,10^, and their simplicity raises the question of why, compared to *Riboviria*, there are so few known examples of viroids and similar agents. Recently, enquiries based on protein similarity have uncovered new Delta-like agents ^2,11^. Likewise, viroids, which lack any protein-coding capacity, are beginning to be surveyed at a larger scale based in part on circular genome maps and the presence of ribozyme-like features. These searches have led to an expanded family of known viroid-like RNAs and a revision of earlier models that their distribution is limited to plants ^12–14^. As such, these studies have already shifted virological paradigms, leaving open the possibility that an even broader category of viroid-like elements are present in living systems that might have been overlooked due to a lack of detectable similarity to known viroids and HDV family members.

The human gut microbiome (hGMB) is experimentally attractive for discovery of novel genetic agents. Indeed, metagenomic and metatranscriptomic ^15^ profiling of the hGMB has yielded new insights into prokaryotic, viral ^16–18^, and plasmid ^19^ ecology. To this end, we developed a reference-free bioinformatic approach (VNom) to identify novel viroid-like elements. We initially applied `VNom` to published Integrative Human Microbiome Project (iHMP) data ^20^ resulting in the identification of a new class of hGMB-colonising RNA agents, which we term ‘Obelisks’. Obelisks form a distinct phylogenetic group restricted to RNA datasets and lack any evident homology to characterised genomes or viromes. Obelisk RNA reads assemble into ∼1000 nt circles, which are predicted to fold into rod-like RNA secondary structures and code for at least one member of a novel “Oblin” protein superfamily. We further found that a subset of Obelisks harbour Obelisk-specific hammerhead ribozyme motifs. Querying 5.4 million public sequencing datasets, we identified 29,959 distinct Obelisks (90 % ID threshold) present across ∼220,000 datasets representing diverse ecosystems beyond the hGMB. Amongst the datasets with clear Obelisk representatives, we identified a definitive Obelisk-Host pair, with *Streptococcus sanguinis* acting as a replicative host. Lastly, we surveyed Obelisks in five published human oral and gut microbiome studies from 472 donors, finding an estimated ∼9.7 % donor prevalence within these datasets, with an apparent anatomy-specific Obelisk distribution.

## Results

### A novel, human microbiome-associated viroid-like RNA

Viroids and Delta viruses are in part typified by their single stranded, circular genomes, both of which are molecular features that can be detected in strand specific RNA-seq. To search for such features in microbiome RNA sequencing (RNA-seq) datasets, we created a bioinformatic tool, `VNom` (see VNom, and Supplementary Figure 2), and applied it to microbiome RNA-seq datasets (see Initial Obelisk identification). In particular, we chose an iHMP human stool dataset ^20^ for its strand-specific RNA-seq, its longitudinal nature (regular sampling over ∼1 year), and its cohort size (104 donors), qualities well suited for identifying persistent hGMB colonists.

We next filtered `VNom`-nominated RNAs to retain contigs with no evident homology to the NCBI BLAST (`nt` or `nr`) databases ^21^ (see Initial Obelisk identification). One class of 15 related (<2 % sequence variation, Supplementary Table 1), 1164 nt RNAs stood out with their extended predicted secondary structure reminiscent of HDV and *Pospiviroidae* (Figure 1b, Supplementary Figures 1a-b and 2b). Owing to a strong predicted rod-like secondary structure, we term this group of RNAs Obelisk-*alpha* (Obelisk-ɑ, “*Obelisk_000001*” in Supplementary Table 1). At 1164 nt in length, the rod-like secondary structure was striking because typical mRNA sequences are not predicted to readily fold in this manner (as evidenced by the efforts required to maximise the degree of “rod-ness” in mRNA vaccines ^22^). Based on open reading frame (ORF) predictions, Obelisk-ɑ has the capacity to code for two proteins (202 and 53 amino acids [aa]). Both open reading frames (ORFs) lack evident nucleotide or protein sequence homology when querying a number of reference databases (NCBI `nt`, `nr`, or `CDD` ^23^, `Pfam` ^24^). Tertiary structure protein alignment yielded similar negative results (see Protein tertiary structure prediction). As such, we chose new names, terming these two proteins Oblin-1 and Oblin-2, respectively. We specifically note that despite some similar characteristics between Obelisk-ɑ and HDV (apparently circular, predicted highly structured RNA genome, and ability to code for at least one ∼200 aa ORF, Supplementary Figure 1a), there is no evident sequence homology at the RNA or protein level or structural homology at the protein level between Obelisks and HDV. In further contrast to HDV, whose large hepatitis Delta antigen (L-HDAg) occurs on one strand of the extended HDV predicted secondary structure (Supplementary Figure 1a), the Obelisk-ɑ Oblin-1 encoding region is largely self-complementary within the open reading frame, forming a ∼300 base pair hairpin making up half of the predicted Obelisk-ɑ RNA secondary structure. Obelisk-ɑ sequences were found to occur in 7 of the 104 iHMP donors (Table 1) with donors A-C exhibiting consistent prevalence for over 200 days (Figure 1d, note: positive donors are renamed for brevity, with original donor alias equivalences in Table 1). Further, Obelisk-ɑ sequences were found to largely cluster together based on donor identity, when grouped by sequence variation (Figure 1e). We noted some co-clustering of sequences between donors (A and E in cluster 3, and D and G in cluster 5); this co-clustering could be explained by either transient prevalence or by library cross contamination, as each minor member of such clusters was both low prevalence (few positive timepoints) and low abundance (low counts in positive timepoints) (see Table 1). Regardless of the source of the relatively rare cross-sample reads, Obelisk-ɑ appears to persist within human donors, with each donor appearing to harbour their own distinct ‘strain.’ Lastly, in companion DNA-seq data from this project, no detectable Obelisk reads are found (Table 2). Taken together, these findings are consistent with Obelisk-ɑ representing an as of yet uncharacterised RNA element with viroid-like features that occurs in human stool, further comprised of subspecies that persist in individual donors over time.

**Figure 1.**
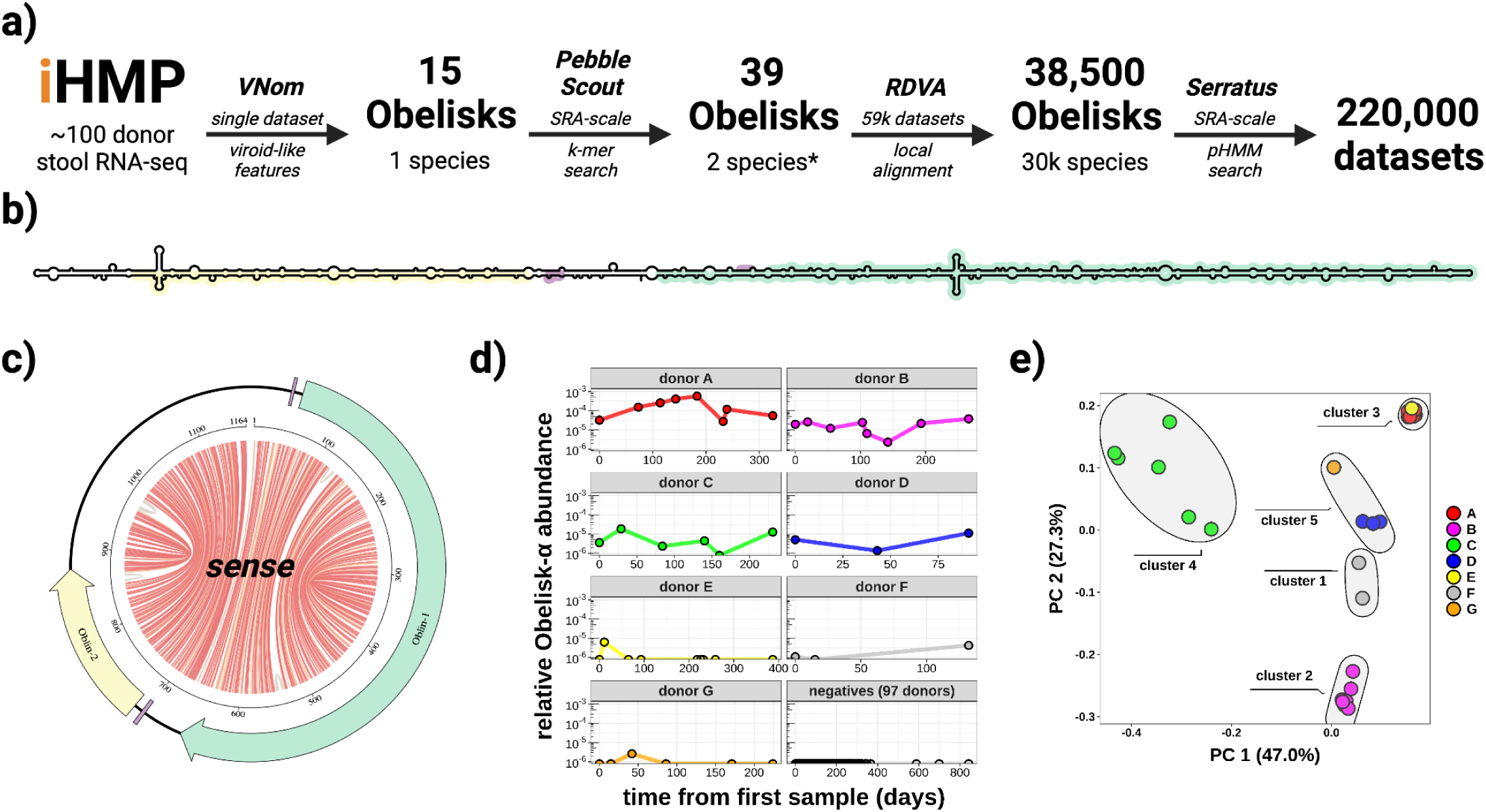
Obelisk *alpha* has a predicted extensive secondary structure and appears to colonise and speciate within the human gut. **a**) overview of the iterative approach taken in Obelisk discovery, (see methods) **b**) schematic of the predicted *sense* consensus secondary structure derived from all non-redundant, 1164 nt Obelisk-ɑs found using SRA-scale k-mer matching (`PebbleScout)`. Predicted open reading frames (ORFs) 1 and 2 (green/yellow), and Shine-Delgarno sequences (purple) shown, **c**) “jupiter” plot of Obelisk-ɑ coloured as in b), chords illustrate predicted basepairs (basepair probabilities grey, 0.1, to red, 1.0) **d**) Obelisk-ɑ relative read abundance for six donors (A-G); sequence data from in *Lloyd-Price et al., 2019* and time in days from first sample. **e**) Principal component analysis of sequence variation seen in Obelisk-ɑ reads in *Lloyd-Price et al., 2019* (the initial iHMP dataset), grouped by k-means clustering with 5 centres, coloured as in d).

**Table 1.** Figure 1d metadata. Donors with at least one Obelisk-*alpha* read from *Lloyd-Price et al. 2019*, with their SRA accession code, alias as used in Figure 1d, disease state, relative (to total reads per sample) Obelisk-ɑ read abundance (see Methods), date of sampling, and original donor numeric ID.

| **SRA**<br>(Lloyd-Price et al. 2019) | **alias** | **clinical**<br>**classification** | **Obelisk-α**<br>**relative cont** | **sample date** | **donor** |
| --- | --- | --- | --- | --- | --- |
| SRR5949245 | donor A | Healthy Donor | 3.22E-05 | 2015-08-31 | d_2077 |
| SRR5950275 | donor A | Healthy Donor | 1.53E-04 | 2015-11-12 | d_2077 |
| SRR5950410 | donor A | Healthy Donor | 2.55E-04 | 2015-12-23 | d_2077 |
| SRR5950468 | donor A | Healthy Donor | 4.06E-04 | 2016-01-21 | d_2077 |
| SRR5950280 | donor A | Healthy Donor | 5.71E-04 | 2016-03-01 | d_2077 |
| SRR5950352 | donor A | Healthy Donor | 2.75E-05 | 2016-04-19 | d_2077 |
| SRR5950308 | donor A | Healthy Donor | 1.19E-04 | 2016-04-26 | d_2077 |
| SRR5950313 | donor A | Healthy Donor | 5.50E-05 | 2016-07-21 | d_2077 |
| SRR5950297 | donor B | Ulcerative Colitis | 1.97E-05 | 2015-11-12 | d_6038 |
| SRR5950395 | donor B | Ulcerative Colitis | 2.62E-05 | 2015-12-01 | d_6038 |
| SRR5950415 | donor B | Ulcerative Colitis | 1.21E-05 | 2016-01-05 | d_6038 |
| SRR5950446 | donor B | Ulcerative Colitis | 2.45E-05 | 2016-02-23 | d_6038 |
| SRR5950283 | donor B | Ulcerative Colitis | 6.46E-06 | 2016-03-01 | d_6038 |
| SRR5950351 | donor B | Ulcerative Colitis | 2.31E-06 | 2016-04-02 | d_6038 |
| SRR5950317 | donor B | Ulcerative Colitis | 2.18E-05 | 2016-05-23 | d_6038 |
| SRR5950257 | donor B | Ulcerative Colitis | 3.78E-05 | 2016-08-04 | d_6038 |
| SRR5949129 | donor C | Crohn's Disease | 3.61E-06 | 2015-04-28 | d_2068 |
| SRR5949216 | donor C | Crohn's Disease | 1.88E-05 | 2015-05-27 | d_2068 |
| SRR5949201 | donor C | Crohn's Disease | 2.34E-06 | 2015-07-21 | d_2068 |
| SRR5950374 | donor C | Crohn's Disease | 4.46E-06 | 2015-09-15 | d_2068 |
| SRR5950426 | donor C | Crohn's Disease | 0 | 2015-10-05 | d_2068 |
| SRR5950333 | donor C | Crohn's Disease | 1.28E-05 | 2015-12-15 | d_2068 |
| SRR5949407 | donor D | Crohn's Disease | 5.09E-06 | 2014-09-16 | d_5001 |
| SRR5949179 | donor D | Crohn's Disease | 1.37E-06 | 2014-10-29 | d_5001 |
| SRR5949326 | donor D | Crohn's Disease | 1.12E-05 | 2014-12-16 | d_5001 |
| SRR5949222 | donor E | Ulcerative Colitis | 0 | 2015-06-15 | d_2071 |
| SRR5949120 | donor E | Ulcerative Colitis | 6.33E-06 | 2015-06-26 | d_2071 |
| SRR5949586 | donor E | Ulcerative Colitis | 0 | 2015-08-19 | d_2071 |
| SRR5950375 | donor E | Ulcerative Colitis | 0 | 2015-09-16 | d_2071 |
| SRR5950383 | donor E | Ulcerative Colitis | 0 | 2016-01-19 | d_2071 |
| SRR5963925 | donor E | Ulcerative Colitis | 0 | 2016-01-27 | d_2071 |
| SRR5950464 | donor E | Ulcerative Colitis | 0 | 2016-02-02 | d_2071 |
| SRR5950281 | donor E | Ulcerative Colitis | 0 | 2016-02-29 | d_2071 |
| SRR5950429 | donor E | Ulcerative Colitis | 0 | 2016-07-06 | d_2071 |
| SRR5949469 | donor F | Crohn's Disease | 1.09E-06 | 2014-08-13 | d_2025 |
| SRR5949284 | donor F | Crohn's Disease | 0 | 2014-08-28 | d_2025 |
| SRR5949607 | donor F | Crohn's Disease | 4.24E-06 | 2014-12-23 | d_2025 |
| SRR5949267 | donor G | Ulcerative Colitis | 0 | 2015-06-23 | d_3034 |
| SRR5949148 | donor G | Ulcerative Colitis | 0 | 2015-07-08 | d_3034 |
| SRR5949435 | donor G | Ulcerative Colitis | 2.58E-06 | 2015-08-04 | d_3034 |
| SRR5950377 | donor G | Ulcerative Colitis | 0 | 2015-09-17 | d_3034 |
| SRR5950332 | donor G | Ulcerative Colitis | 0 | 2015-12-11 | d_3034 |
| SRR5950460 | donor G | Ulcerative Colitis | 0 | 2016-02-02 | d_3034 |

**Table 2.** Obelisk read counts from paired metagenomic (DNA) and metatranscriptomic (RNA) samples. Paired metagenomic DNA and metatranscriptomic RNA `Kraken2` read counts (`Bracken`-corrected, see methods) for *Lloyd-Price et al. 2019* ^20^ and *Maghini and Dvorak et al. 2023* ^79^ for Obelisks -ɑ and -β. Means taken over donors (aliased as in Table 6) and total read counts also reported. Note that no reads mapping to either Obelisk were found in the DNA datasets, suggesting an RNA-only Obelisk lifestyle. For full `Kraken2` and `Bracken` outputs, see Data Availability.

| study | donor | mean<br>$\alpha$ RNA | sdv $\alpha$<br>RNA | mean<br>$\beta$ RNA | sdv $\beta$<br>RNA | mean<br>RNA<br>reads | sdv RNA<br>reads | mean<br>$\alpha$ DNA | sdv $\alpha$<br>DNA | mean<br>$\beta$ DNA | sdv $\beta$<br>DNA | mean<br>DNA<br>reads | sdv DNA<br>reads |
| --- | --- | --- | --- | --- | --- | --- | --- | --- | --- | --- | --- | --- | --- |
| Lloyd-Price *et al.* 2019 | A | 2310 | 3180 | 1 | 1 | 13372847 | 8928775 | 0 | 0 | 0 | 0 | 12434953 | 7135963 |
| Lloyd-Price *et al.* 2019 | B | 120 | 75 | 0 | 0 | 6471339 | 1941900 | 0 | 0 | 0 | 0 | 8624205 | 2706287 |
| Lloyd-Price *et al.* 2019 | C | 59 | 85 | 0 | 0 | 7465073 | 3571490 | 0 | 0 | 0 | 0 | 7861229 | 2987857 |
| Lloyd-Price *et al.* 2019 | D | 35 | 36 | 0 | 0 | 5991299 | 1931426 | 0 | 0 | 0 | 0 | 11930216 | 3223559 |
| Lloyd-Price *et al.* 2019 | E | 5 | 13 | 0 | 0 | 10205726 | 4505426 | 0 | 0 | 0 | 0 | 10182743 | 4150798 |
| Lloyd-Price *et al.* 2019 | F | 14 | 6 | 0 | 0 | 7261820 | 2583126 | 0 | 0 | 0 | 0 | 9293180 | 3041056 |
| Lloyd-Price *et al.* 2019 | G | 3 | 4 | 0 | 0 | 7544276 | 3692007 | 0 | 0 | 0 | 0 | 8342623 | 2564388 |
| Maghini and Dvorak *et al.* 2023 | 1 | 8080 | 5500 | 319 | 246 | 24534988 | 11197080 | 0 | 0 | 0 | 0 | 59395503 | 52132729 |
| Maghini and Dvorak *et al.* 2023 | 2 | 0 | 0 | 0 | 0 | 17277835 | 6182452 | 0 | 0 | 0 | 0 | 63203793 | 31663431 |
| Maghini and Dvorak *et al.* 2023 | 3 | 0 | 0 | 0 | 0 | 14125535 | 9993287 | 0 | 0 | 0 | 0 | 35879627 | 10380606 |
| Maghini and Dvorak *et al.* 2023 | 4 | 0 | 0 | 0 | 0 | 15983644 | 5232547 | 0 | 0 | 0 | 0 | 19114616 | 10718757 |
| Maghini and Dvorak *et al.* 2023 | 5 | 24 | 42 | 0 | 0 | 18237767 | 5281905 | 0 | 0 | 0 | 0 | 38104238 | 18496583 |
| Maghini and Dvorak *et al.* 2023 | 6 | 0 | 0 | 0 | 0 | 19476656 | 3667861 | 0 | 0 | 0 | 0 | 44924227 | 19547694 |
| Maghini and Dvorak *et al.* 2023 | 7 | 0 | 0 | 0 | 0 | 18639827 | 6428812 | 0 | 0 | 0 | 0 | 75407696 | 72530680 |
| Maghini and Dvorak *et al.* 2023 | 8 | 4 | 6 | 0 | 0 | 18449872 | 6842782 | 0 | 0 | 0 | 0 | 27950881 | 10870547 |
| Maghini and Dvorak *et al.* 2023 | 9 | 1 | 1 | 0 | 0 | 15639556 | 9328742 | 0 | 0 | 0 | 0 | 42868581 | 32749586 |
| Maghini and Dvorak *et al.* 2023 | 10 | 12417 | 1740 | 1 | 2 | 13021384 | 4204129 | 0 | 0 | 0 | 0 | 20243492 | 8732203 |

### Public data are replete with Obelisk-like elements

Using Obelisk-ɑ as a starting point, 21 additional full-length examples of Obelisk-ɑ (<4 % sequence variation, Supplementary Table 1) were found in 7 datasets using a k-mer search (`PebbleScout` ^25^) of ∼3.2 million “metagenomic” annotated sequence read archive (SRA) datasets. All 7 datasets were human-derived metatranscriptome (metagenomic RNA) BioProjects (Table 3, see Obelisk homologue detection in additional public data); 0 sequences were found in metagenomic DNA samples. The repeated finding of Obelisk-ɑ in disparate BioProjects supported the notion that Obelisk-ɑ is a *bona fide* biological entity. Based on the prevalence of Obelisk-ɑ in these human microbiome transcriptome datasets (Table 3), we investigated the possibility that additional Obelisks might be present in such data (as identified by both `VNom` and Oblin-1 protein similarity). This search ultimately lead to the discovery of Obelisk-β (“*Obelisk_000002*” in Supplementary Table 1), a 1182 nt, likely hGMB-resident, Obelisk-like RNA with similar characteristics to Obelisk-ɑ (circular assembly map, rod-like predicted secondary structure, absence in paired DNA sequence) and low-but-evident protein sequence similarity to Oblin-1 (∼38 % protein similarity and pairwise mean `BLASTp` E-value: 5.2×10^-14^). Thus, both Obelisks appear to be Oblin-1-encoding elements. Analysis of the Oblin-2 homology at this stage was limited by the short size of the proteins – nonetheless both Obelisk-ɑ and Obelisk-β encode second proteins of ∼50 amino acids rich in helix-forming residues (Supplementary Figure S3a/c). Next, utilising the uniqueness of the Obelisk-ɑ/β Oblins-1 and -2 as Obelisk-specific hallmark sequences, we searched over 12 trillion contigs in the `RNA deep virome assemblage` (`RDVA`), a database of assembled metatranscriptomes ^13,26^ (see Obelisk homologue detection in additional public data), yielding over 38,500 Oblin-encoding RNA assemblies. Following this search, the smaller Obelisk-ɑ/β proteins were determined to likely be Oblin-2 homologues (∼31 % protein similarity and mean `BLASTp` E-value against the Oblin-2 consensus sequence: 2.5×10^-6^). Ultimately, by insisting on evidence of apparent circularity, we created a stringent subset of 7,202 clustered Obelisks (1,744 clusters at 80 % nucleotide identity) as a conservative database for future studies (Supplementary Table 1). Building from these `RDVA` hits, we then queried ∼5.4 million SRA datasets for distant Oblin-1 and -2 homology using `Serratus` ^2^ (applying an inclusion threshold from earlier `Serratus` projects, see Serratus), yielding over 220,000 putatively Obelisk-positive datasets. From these datasets, we followed up on the 4,505 datasets with confident Oblin-1 hits (see Serratus). These searches suggest that Obelisk-like elements are found globally (Figure 3c), with a possible bias in reported datasets towards mammalian microbiome-related origins (Figure 3b), and represent a distinct, diverse group of phylogenetically related RNA-based elements (Figure 3a).

**Figure 2.**
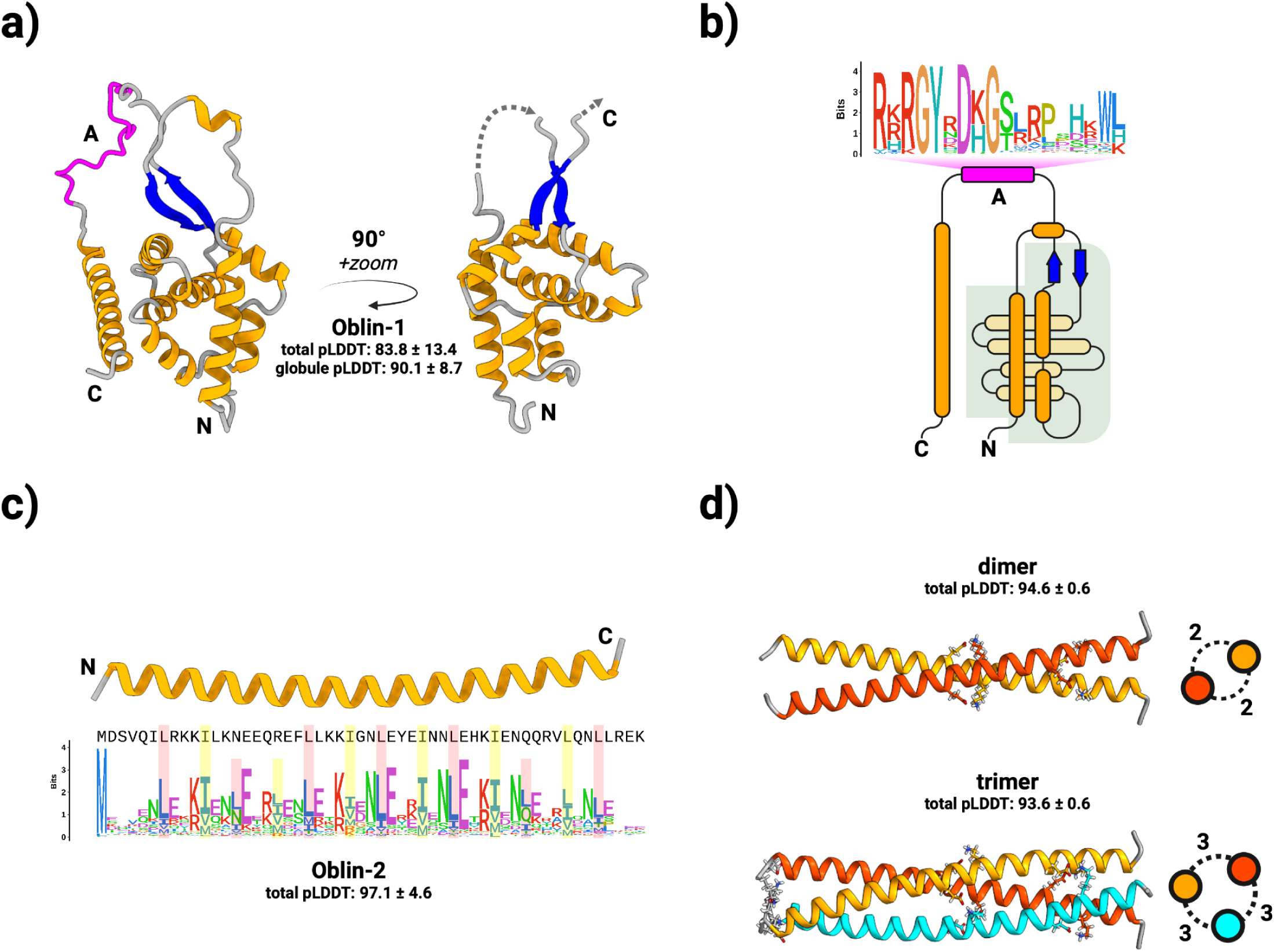
Obelisks encode putatively well-folded proteins. **a**) Obelisk open reading frame 1 (Oblin-1) is predicted (total mean-pLDDT ± SD = 83.8 ± 13.4, see methods) to fold into a stereotyped N-terminal “globule” formed of a three alpha helix (orange) bundle partially wrapping around an orthogonal four helix bundle, capped with a beta sheet “clasp” (blue, globule mean-pLDDT = 90.1 ± 8.7), joined by an intervening region harbouring the conserved *domain-A* (magenta) with no predicted tertiary structure, to an arbitrarily placed C-terminal alpha helix. “Globule” emphasised on the right. **b**) a to-scale (secondary structure) topological representation of Oblin-1 with the “globule” shaded in grey, and the *domain-A* emphasised with this bit-score sequence logo (see methods). **c**) Obelisk Oblin-2 is confidently predicted (mean-pLDDT = 97.1 ± 4.6 ) to fold into an alpha helix which appears to be a leucine zipper. Sequence logo of an “i+7” leucine spacing emphasised in red, with hydrophobic “d” position residues emphasised in yellow (expanded in Supplementary Figure 4b). **d**) homo-multimer predictions of Obelisk-*alpha* Oblin-2. **top:** dimer (mean-pLDDT = 94.6 ± 0.6), **bottom:** trimer (mean-pLDDT = 93.6 ± 0.6). Side-on representations of homomultimers shown with numbers of inter-helix salt-bridges (see Supplementary Figure 5).

**Figure 3.**
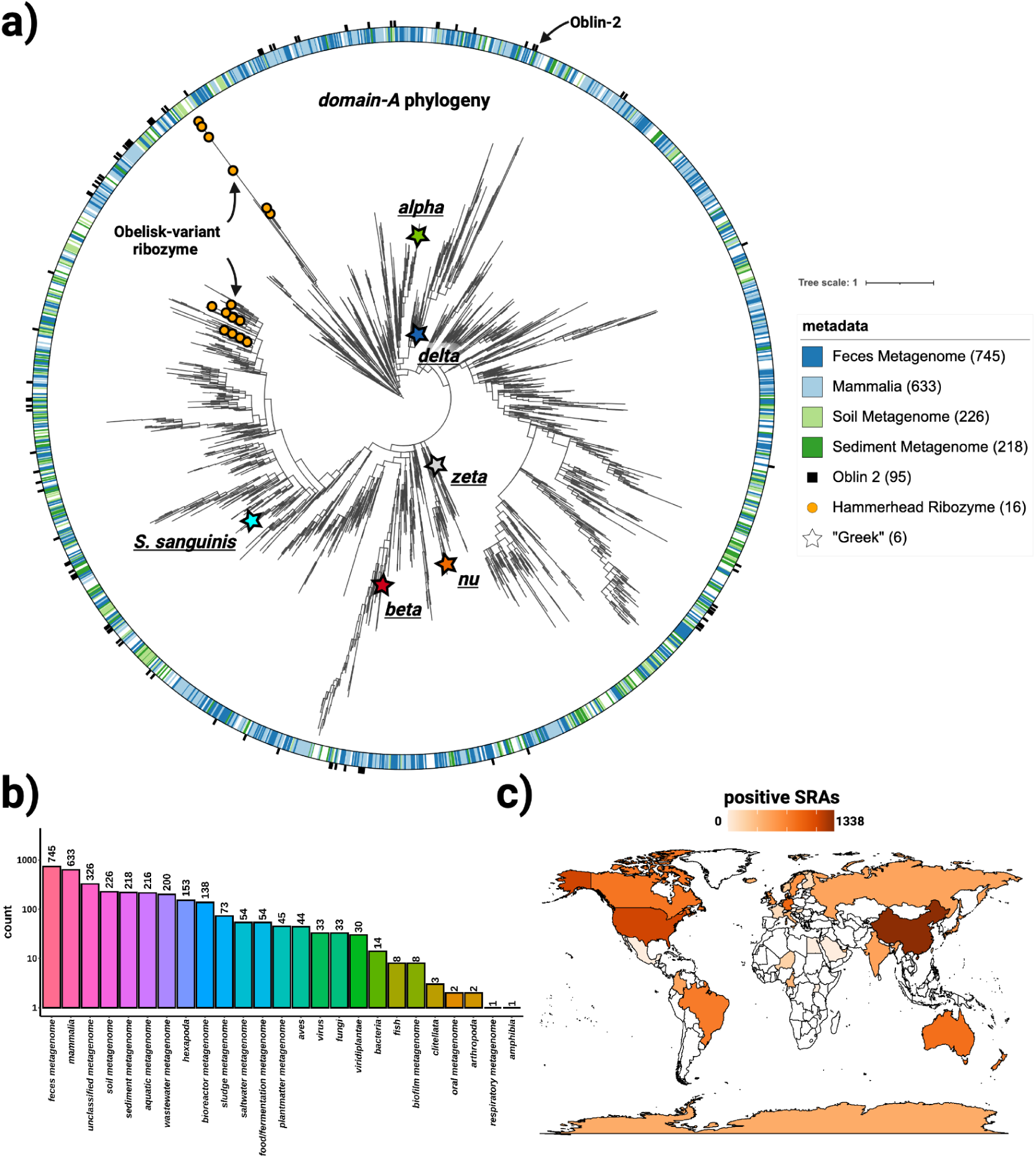
Obelisks form their own globally distributed phylogenetic group. **a**) a maximum likelihood, midpoint-rooted, phylogenetic tree (see methods) constructed from a non-redundant set of 3265 `Serratus` and `RDVA` *domain-A* sequences, with `RDVA` genomes positive for Obelisk-variant self cleaving Hammerhead Type III ribozymes illustrated as orange circles on leaves, and the top four known classes of SRA “host” metadata depicted as the colour band (see legend), and with per-`RDVA`-genome co-occurrence of Oblin-2 (based on `blastp` hits against the Oblin-2 consensus) illustrated as the outer ring (black studs). Leaves that correspond to *domain-A* sequences from Figure 4 are illustrated with stars. **b**) Counts of non de-replicated `SRA` datasets used to construct a) sorted by their “host” metadata; we note that “host” metadata likely fails to account other organisms’ genetic material that was sequences alongside the “host” (e.g. signals from these hosts’ microbiomes maybe be detected in tandem). **c**) Counts of non de-replicated `SRA` datasets used to construct a) arranged by sample geolocation (where known) illustrated on a world map (darker orange = more `SRA` datasets contributed to a)). We note that `SRA` counts are not expected to correlate with true geo-/ecological prevalence, but are still indicative of global presence.

**Table 3.** varied human stool metatranscriptomes are positive for Obelisk-*alpha*. Obelisk-*alpha* positive bioProjects identified by `PebbleScout` (see methods)

| bioProject | human stool study details |
| --- | --- |
| PRJNA398089 | longitudinal - part of iHMP: where Obelisk-alpha was first identified |
| PRJNA940499 | testing stool storage conditions: where Obelisk-beta was first identified |
| PRJNA407499 | longitudinal - typhoid vaccine challenge trial |
| PRJNA354235 | longitudinal - part of the Health Professionals Follow-up Study |
| PRJNA600008 | secondary bile acid intestinal inflammation |
| PRJNA541981 | pre immunotherapy treatment in melanoma patients |
| PRJNA338184 | longitudinal - enterotoxigenic Escherichia coli challenge |
| PRJNA306874 | longitudinal - part of Inflammatory Bowel Disease Multi-omics Data (IBDMDB) |
| PRJNA389280 | longitudinal - part of iHMP |

### An oral commensal bacterium, *Streptococcus sanguinis,* serves as one Obelisk host

The task of identifying specific host-agent pairings from metagenomic data presented a number of challenges. Most samples with Obelisk homologues that were retrieved from the various searches were from metatranscriptomic samples derived from complex mixtures such as highly biodiverse microbiome and waste water samples (Figure 3b). As such, the potential host(s) of Obelisk elements were not immediately clear. While correlation and co-occurrence based methods for inferring potential hosts are possible ^3,4,16^, concerns about their statistical validity and interpretability ^27–29^ motivated a more direct strategy for Obelisk host identification. Consequently, we combed the `Serratus` results for Obelisk-like elements found in limited-complexity samples, such as defined monoculture and/or co-cultures. This search yielded a set of independent sequencing datasets from *Streptococcus sanguinis* (strain SK36), a commensal bacterium of the healthy human oral microbiome ^30^. Several RNA-seq datasets (Table 4) from *S. sanguinis* strain SK36 contained an Oblin-1 coding Obelisk-like sequence (see *Streptococcus sanguinis* bioinformatics). These datasets evidenced a well-defined RNA element which we refer to as “Obelisk-*S.s*” (“*Obelisk_000003*” in Supplementary Table 1). This RNA has the hallmark features of an Obelisk: a characteristic length (1137 nt) circular assembly with an obelisk-shaped predicted RNA secondary structure; genome similarity to Obelisks-ɑ and -β (41 and 35 % nucleotide identity, respectively) and an Oblin-1 homologue (ɑ and β: 33 % protein similarity, and mean pairwise E-values of 5.2×10^-5^ and 4.5×10^-7^, respectively). Unlike the other two Obelisks, however, it lacks a predicted Oblin-2 homologue (Supplementary Figure S2a/d). Overall, based on sequence homology, the predicted genomic secondary structure, the Oblin-1 tertiary structure, and the Obelisk-characteristic Oblin-1 self-complementarity (Supplementary Figure S2d), this RNA element is a *bona fide* Obelisk. Further, the robust co-occurrence of *S. sanguinis* SK36 with Obelisk RNA-seq reads (Table 4), positions *S. sanguinis* SK36 as a model system for future Obelisk characterisation.

**Table 4.**
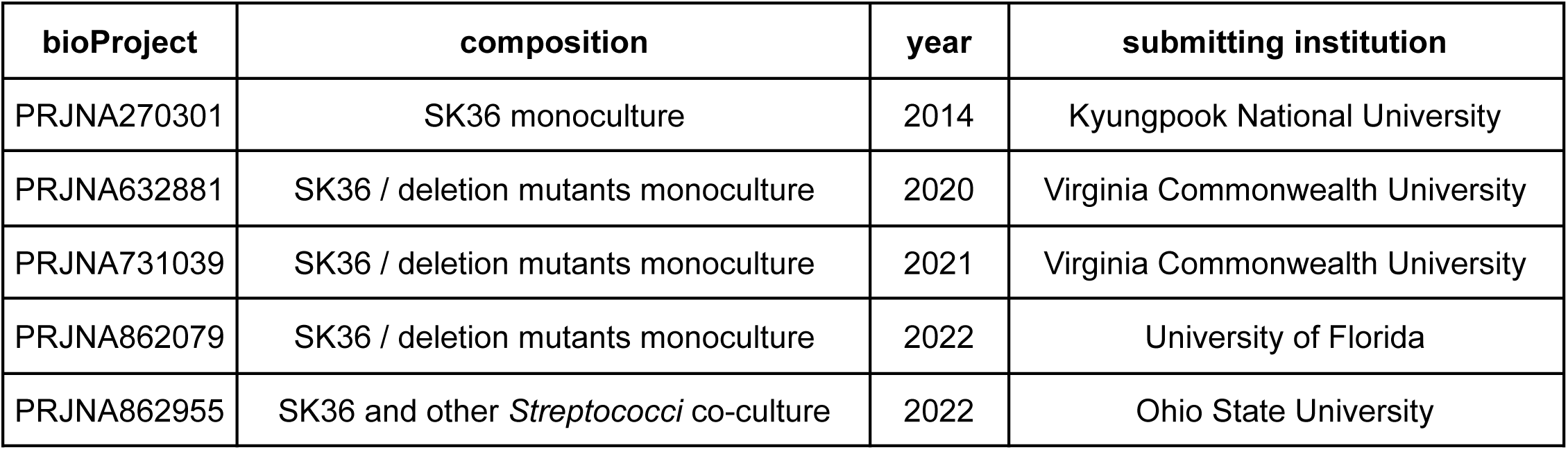
multiple *Streptococcus sanguinis* SK36 datasets contain Obelisk reads. A curated set of *Streptococcus sanguinis* SK36 monoculture or low complexity bioProjects identified by `scanRabbit` (see methods) that contain Obelisk reads produced from differing institutions over differing years and thus suggestive of an *S. sanguinis* SK36 - Obelisk relationship.

### Structural prediction indicates a novel globular domain characteristic of Oblin-1 proteins

Due to the lack of obvious protein sequence homology in existing, non-Obelisk databases, we performed protein tertiary structure predictions in an attempt to identify both shared predicted structural elements, and homology through tertiary structure similarity searches. Owing to Oblin-1 and -2’s previously unrecorded nature and apparent monophyly, we avoided automated multiple sequence alignment construction during conventional tertiary structure prediction using `ColabFold` (an implementation of `AlphaFold2`) ^31,32^ and instead opted for custom `RDVA` alignments (see Protein tertiary structure prediction). This yielded a folding prediction of Oblin-1 (mean per-residue confidence estimate, μ-pLDDT ± standard deviation, of 83.8 ± 13.4, where 70-90 pLDDT values are “*a generally good backbone prediction*” ^33^ and higher is better) with a more confidently predicted N-terminal “globule” (μ-pLDDT of 90.1 ± 8.7, Figure 2a). `ColabFold` was not able to confidently place the two flanking backbones between the first and last predicted alpha helices and the rest of the Oblin-1 sequence (Supplementary Figure 4a), and owing to heterogeneity in the last alpha helix’s placement across predictions (see Data Availability), the “globule” was further focused on. The “globule” was predicted to form a consistent fold (Figure 4): a three alpha helix bundle (two smaller alpha helices co-axially aligned along the larger alpha helix) partially wrapping over a semi-orthogonal four alpha helix bundle - all bookended with a two strand beta sheet “clasp” (Figure 2b). Interestingly, no confident fold was predicted for the largest conserved region in Oblin-1 (Supplementary Figure 4), termed *domain-A*, (Figure 2a-b - magenta). Suggestive of an anion binding function, this 18 amino acid stretch is enriched for positively charged residues (arginine, histidine, and lysine) with the Obelisk-ɑ *domain-A* containing five arginines, three histidines, and a lysine residue (50 % of *domain-A*, Figure 2b and Protein homology bioinformatics). Additionally, a “`GYxDxG`” motif appears prominently in *domain-A*. If Oblin-1, represents a new class of RNA binding protein, `ColabFold` may miss the fold of *domain-A* due to the absence of its client RNA ligand.

**Figure 4.**
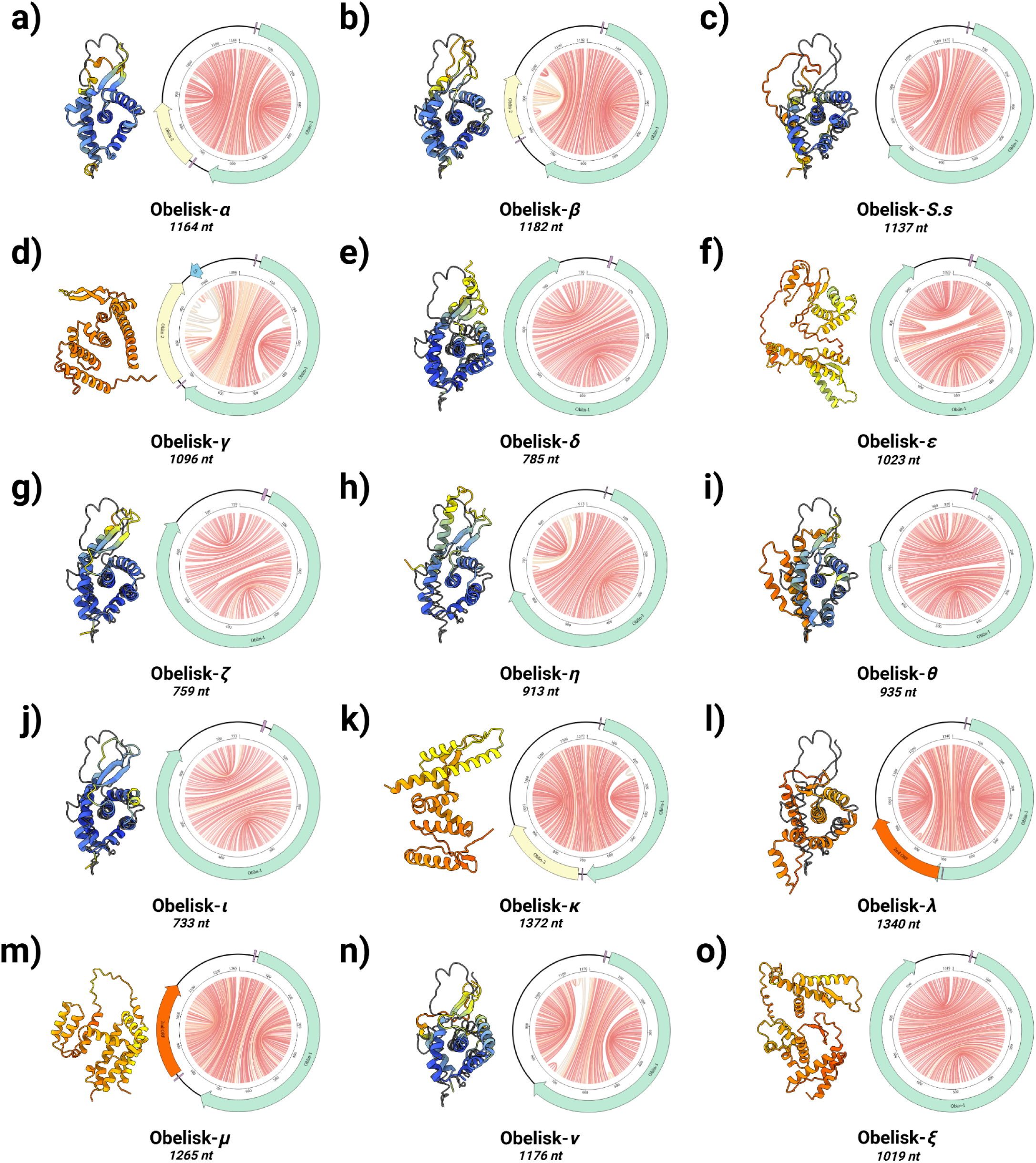
Obelisks form a self-consistent set. Predicted Obelisk secondary structures depicted as “jupiter” plots where chords represent predicted basepairs (coloured by basepair probability from 0, grey, to 1, red, see methods) with predicted open reading frames (ORFs, preceded by predicted Shine-Delgarno sequences, purple) depicted: Oblin-1 (green), Oblin-2 (yellow, based on `blastp` hits against the Oblin-2 consensus), and “2ndORF” (orange). Obelisk-ɣ’s suggested CRISPR spacer match illustrated in light blue. `ColabFold` predictions of Oblin-1 tertiary “globule” structures built with *ad hoc* multiple sequence alignment (MSA) construction (coloured cartoons) superimposed over the `RDVA`-derived MSA prediction for Obelisk-α where possible (black line, Figure 2a, see methods). Prediction confidence (pLDDT) shown as cartoon colouring as in Supplementary Figure 3. Greek letter key: α : alpha, β : beta, ɣ : gamma, δ : delta, ε : epsilon, ζ : zeta, η : eta, θ : theta, ι : iota, κ : kappa, λ : lambda, μ : mu, ν : nu, and ξ : xi.

### Oblin-2 modelling suggests a leucine zipper alpha helix

Oblin-2 modelling with `ColabFold` resulted in a high-confidence prediction (μ-pLDDT of 97.1 ± 4.6) that this protein forms a solitary alpha helix (Figure 2c). In the `RDVA` consensus, the Oblin-2 alpha helix consists of a leucine zipper motif (see Protein conservation and phylogenetics), with the characteristic “i+7” spacing of leucines at the “a” position; another hydrophobic residue (leucine or isoleucine) with “i+7” spacing at the “d” position; and complementary charged residues (glutamic acid and lysine or arginine) at the “e” and “g” positions, respectively ^34^ (Supplementary Figure 4b). Based on μ-pLDDT, `ColabFold` predicts that Oblin-2 might be able to homo-multimerize as a dimer (μ-pLDDT of 94.6 ± 0.6), or a trimer (μ-pLDDT of 93.6 ± 0.6) with a coiled-coil forming with 2 or 3 inter-helix salt bridges per helix pair, respectively (Figure 2d, and Supplementary Figure 5a-b). Although conceivable, a higher order Oblin-2 homo-tetramer is less well supported by `ColabFold` (μ-pLDDT of 65.3 ± 7.9, Supplementary Figure 5c). Leucine zippers typically act as multimerization motifs that bring together other protein domains such as the DNA-binding basic leucine zipper domain (bZIP) ^35^. Oblin-2 does not appear to include any other sequence motifs (*e.g*. a non-zipper poly-basic patch similar to bZIP proteins), suggesting potential function as a homo-multimer, or as a binding partner to other host leucine zippers.

### A subtype of Obelisks bear ribozyme signatures of a viroid-like replication mechanism

Viroids of the family *Avsunviroidae* and HDV code for self-cleaving ribozymes used in their respective replicative cycles ^6,8^ (Supplementary Figure 1a,c), and previous bioinformatic studies have found self-cleaving ribozymes in candidate viroid-like genomes ^12–14^. Upon querying for Hammerhead type-III ribozyme-coding Obelisks, we identified 23 initial hits and noticed that these ribozymes slightly differed from the reference covariance model (`Rfam: RF00008`). Therefore, we constructed an “Obelisk-variant Hammerhead type-III” ribozyme (ObV-HHR3) covariance model (Supplementary Figure 6b, see RNA homology bioinformatics), yielding 339 total Obelisks containing HHRs in the `RDVA` set with stringent similarity (35 clustered at 80 % identity in Supplementary Table 1 - “ObV-HHR3” column). These “HHR-Obelisks” are similarly rod-shaped, ∼1 kb in length, and code for diverged Oblin-1 proteins (20.6 % identity and 31.7 % similarity to the Obelisk-ɑ Oblin-1) that are similarly largely self-complementary (Supplementary Figure 6a), do not code for Oblin-2, but do include an unrelated “smaller ORF.” Additionally, some Obelisks appear to include a bidirectional pair of ObV-HHR3 ribozymes (Supplementary Figure 6a), a feature used by *Avsunviroidae*, HDV, and ambiviruses for their rolling-circle replicative cycles. For the subset of ObV-HHR3 ribozyme-containing Obelisks, `ColabFold` predicts a “globule” fold (total μ-pLDDT of 76.8 ± 20.1, and “globule” μ-pLDDT of 88.3 ± 8.6, Supplementary Figure 6c, Supplementary Figure 7), that is similar to the non-HHR Oblin-1 model but with additional specific tertiary structure features. Namely, the beta-sheet “clasp” region is expanded by an extra sheet as well as some small alpha helices, and the C-terminal alpha helix is predicted to be shorter (Supplementary Figure 6d). Additionally, the *domain-A* region appears to be diverged in the ObV-HHR3 class, yet still exhibits the positive residue skew as well as the “`GYxDxG`” protein motif also found in non-HHR-Obelisks (Supplementary Figure 6d). These subset-specific features, and the correlation with HHR co-occurrence, suggest that at least HRR-Obelisks may replicate via a viroid-like mechanism, with Oblin-1 and/or Oblin-2 as potential cofactors.

### A phylogeny of Oblin-1’s *domain-A* provides evidence for in-family evolution and places ribozyme-baring Obelisks in distinct clades

Following the `RDVA` and `Serratus` searches, an initial Obelisk phylogeny spanning diverse sampling sites (Figure 3b) from around the globe (Figure 3c) was constructed using *domain-A* as a marker sequence (see Protein homology bioinformatics and Protein conservation and phylogenetics, Figure 3a). This *domain-A* phylogeny was sufficient to partially explain the distribution of ObV-HHR3-bearing Obelisks, which segregate tightly into two clades (Figure 3a - orange circles), implying both an evolutionary relationship between Obelisk genome processing and *domain-A*, as well as two different evolutionary paths for *domain-A* ‘speciation’ within the presence of ObV-HHR3. Additionally, this phylogeny indicates that the human microbiome-associated Obelisks (Figure 3a - stars) are widely distributed, implying a complex intersection between human and Obelisk biology. However, the co-occurrence of Oblin-2 (Figure 3a - black studs), and the sampling site of origin (Figure 3a - coloured band), are not adequately explained by this *domain-A* phylogeny, suggesting either multiple gains or losses of such features over the course of Obelisk evolution or recombination events that would confound the construction of a simple tree.

### Absence of captured Obelisk matches among available CRISPR spacer datasets

Searches through CRISPR spacer databases offer an opportunity to deduce past associations between specific mobile genetic elements and potential cellular prokaryotic hosts ^4,14^. We applied a conservative k-mer matching approach (see Obelisk spacer analysis) to gauge the extent to which Obelisks appear to be sampled by the CRISPR spacer arrays, using a dataset of 29,857,318 spacers predicted by the Joint Genome Institute’s (JGI’s) `IMG/M` database ^36^. Ultimately, only one spacer locus out of ∼140,000 initially mapping spacers confidently mapped to an >1000 nt Obelisk-like contig which we term Obelisk-“gamma” (Obelisk-ɣ, Supplementary Figure 8, “*Obelisk_000004*” in Supplementary Table 1). This mapping could suggest that Obelisk-ɣ has previously infected the Alphaproteobacterium *Bombella mellum*, however, the Obelisk that this spacer maps to deviates from the “rod-like” nature seen in other Obelisks (Figure 4 - “jupiter” plots), suggestive of a chimeric misassembly. While Obelisk-ɣ does resemble other Obelisks (1096 nt, mostly rod-shaped, and contains both Oblin-1 and Oblin-2 homologues - Supplementary Figure 8c), the assembly appears to contain an unpaired, “frayed” end. The coincidence of the spacer mapping to the “frayed” end, and the fact that only *one* mapping was found (out of ∼39,000 `RDVA` Obelisks) casts some doubt on the validity of this mapping. As such, using this spacer mapping approach, we have no evidence to date that CRISPR systems interact with Obelisks, but that if they do, these interactions appear to be rare events or make use of a CRISPR system that has not been appended to the `IMG/M` database. Alternatively, the surveyed Obelisks, and the methods used to identify them, might have serendipitously been biassed against identifying CRISPR-interacting Obelisks.

### Obelisks are prevalent in tested human microbiomes

Next, we sought to roughly estimate the prevalence of Obelisks in human gut and oral microbiomes by searching five datasets (three gut, two oral, Table 5) spanning 472 human donors primarily from North America (due to representational bias on the `SRA`). 25 donors (5.3 %) were identified as positive for Obelisks -ɑ, -β, or -*S.s*, and a further 21 donors (4.4 %) appeared to be positive for novel Obelisks (Supplementary Figure 9), for a total of 9.7 % Obelisk-positivity (see Surveying for Obelisks in human data). Upon separating by microbiome source, 6.6 % (29 donors) of gut microbiome, and 53 % (17 donors) of oral microbiome samples contained Obelisks. These data therefore implicate the oral microbiome as a reservoir of Obelisks with more than half of the donors positive for such elements, though this could also be explained by an idiosyncrasy of the major oral dataset (*Belstrøm and Constancias et al. 2021* ^112^) that contributes to this count. Ultimately, 11 new, distinct, full-length Obelisks were identified upon examining the Obelisk-positive donors without Obelisk -ɑ, -β, or -*S.s* homology - which we name “delta” through “xi” (see Surveying for Obelisks in human data, Figure 4, “*Obelisk_000005*” through “*Obelisk_000015*” in Supplementary Table 1). Obelisks “alpha,” “beta,” “epsilon,” “zeta,” and “eta” were restricted to gut microbiome samples (Obelisk-ε was found in one oral sample), whereas Obelisks “*S. sanguinis*,” and “theta” through “xi” were primarily orally restricted (Obelisk-*S.s* was found in one stool sample) - indicating an anatomical specificity of Obelisks despite the oral-gastric connection. These studies used different library preparation strategies (Table 5) and show varying Obelisk sensitivity as a function of read depth (Supplementary Figure 9 - scale bars), consistent with the technical expectation that not all metatranscriptomic sequencing workflows would be equally good at detecting Obelisks. This raises the question of a potential technological blind-spot to these (and similar) elements with some protocols. In any case, the observed values certainly represent a lower bound, and these data point to Obelisks being a non-negligible member of the tested adult oral and gut microbiomes. By their public nature, these datasets lack complete donor medical metadata; this lack and the relatively small sample size leave the investigation of correlations between Obelisk prevalence (and abundance) and the health of human hosts for future studies.

**Table 5.** human oral and stool metatranscriptome datasets queried for Obelisks. Human metatranscriptomic datasets used for Supplementary Figure 9 with sampled niche and library preparation method indicated.

| bioProject | study | niche | RNA-seq library method |
| --- | --- | --- | --- |
| PRJNA398089 | Lloyd-Price *et al.* 2019 <sup>20</sup> | stool | Modified RNAtag-seq |
| PRJNA940499 | Maghini and Dvorak *et al.* 2023 <sup>79</sup> | stool | Illumina stranded total rna prep |
| PRJNA812699 | Jacobs *et al.* 2023 <sup>112</sup> | stool | Viomega |
| PRJNA678453 | Belstrøm and Constancias *et al.* 2021 <sup>37</sup> | oral | Illumina TruSeq Stranded mRNA |
| PRJNA917314 | Tong *et al.* 2023 <sup>113</sup> | oral | NEBNext Small RNA-seq |

## Discussion

The RNA viroid/sub-viral component of the biosphere is beginning to be estimated ^12–14^, but sequence-matching-based strategies, though potent for RNA viral discovery ^1–5^, are blind to previously unnoticed classes of agents. Here, we applied a generic molecular-feature-focused search strategy (VNom) to identify viroid-like RNAs in public RNA-seq datasets. We ultimately focused on a large monophyletic group of viroid-like elements that we term Obelisks. A single clear Obelisk-host pairing (*S. Sanguinis* SK36 - Obelisk-*S.s*) indicates that Obelisks can be a component of bacterial cells; while we don’t know the “hosts” of other Obelisks, it is reasonable to assume that at least a fraction may be present in bacteria.

Obelisk genomes are predicted to fold into conspicuous “rod-like” secondary structures, with a largely self-complementary and conserved Oblin-1 ORF that accounts for at least half of the circular sequence assembly. Oblin-1 itself is predicted to fold into a stereotyped “globule” tertiary structure (Figure 4) with its most conserved motif, *domain-A*, lacking a confident tertiary structure prediction (Figure 2a). Furthermore, the presence of a subset of hammerhead ribozyme-bearing Obelisks with distinct Oblin-1 features (Supplementary Figure 6), and an interplay with *domain-A* evolution (Figure 3a) suggests an Oblin-ribozyme functional relationship, perhaps in viroid-like rolling-circle or rolling hairpin ^37^ replication. We note that conservative ribozyme detection thresholds were used in this work, leaving open the possibility that a larger diversity of ribozymes could be present in the Obelisks, including potentially novel self-cleaving ribozymes. As such, the exact interplay between Obelisk genome processing (*via* ribozymes) and Oblins-1, -2, and others is currently unknown.

Obelisks appear to be globally distributed (Figure 3c) and are a constituent member of the human oral and gut microbiomes, occurring in ∼10 % of human donors in five assayed human metatranscriptomic studies (Table 5, Supplementary Figure 9). Of particular interest, we note one oral microbiome study showing a ∼50 % Obelisk prevalence (Supplementary Figure 9d). We also note that observed Obelisk prevalence is likely to be quite dependent on the population in question, sampling scheme, type and depth of sequencing, and other features. Lastly, a specific Obelisk strain, Obelisk-ɑ, appears to persist and speciate within microbiomes of human donors (Figure 1d-e). The prevalence and apparent novelty of these elements implies more is yet to be learned about their interplay with microbial and human life.

Constructing a full Obelisk phylogenetic tree with explanatory power proved difficult (Figure 3a). This is likely due to several factors including the fact that Obelisks appear to be under selection for a highly basepaired genomic coding region that must also code for stereotyped protein fold (Figure 4). Classical phylogenetic tools cannot account for evolutionary signals from non-position-independent RNA secondary structure constraints ^38^, consistent with the complexities in estimating trees from such families ^39^. Further, recent advances in protein tertiary structure prediction may now allow for protein structure based phylogenetic reconstruction that may be tolerant of greater sequence divergence ^40,41^. As such, definitive phylogenetic work on Obelisks might benefit from future tools that incorporate both evolutionary signals from RNA secondary structure conservation, and from structural alignment of predicted Oblin-1 “globule” tertiary structures. Lastly, the `Serratus` approach taken for large-scale Obelisk discovery was run using homology models built from sequences initially homologous to Obelisk-ɑ (see Protein homology bioinformatics), and thresholds derived from RNA viral discovery campaigns ^2^, so while a mammalian sample-origin bias is seen (Figure 3b), this could be explained by an auto-correlation based on the mammalian origin of Obelisk-ɑ, potentially confounded by the choice of RNA viral discovery threshold. Due to this aforementioned bias, as well as a lack of a systematised method for discovery, it should be noted that the breadth of Obelisk diversity reported in this study could be an underestimate. Further, while we focused on Obelisks, their prevalence and diversity suggest that similar, unrelated viroid-like RNAs are likely widespread and waiting to be discovered in public sequencing data.

The observation that distinct subsets of Obelisks appear to occur in human oral versus gastric sites, an anatomic specificity that mirrors the site-specificity of human microbiomes ^42^ (Supplementary Figure 9), supports the notion that Obelisks might include colonists of said human microbiomes. Building on this, donor-specific factors such as diet or lifestyles therefore likely play a role in Obelisk (re-)colonisation and retention. Further, given that *Streptococcus sanguinis* is a commensal of the healthy human oral microbiome ^43^, but also a causative agent of bacterial endocarditis ^44^, study of the implied *S. sanguinis*-Obelisk-*S.s* relationship might begin to reveal the relevance of Obelisks to the natural oral niche and potentially to human health, as well as offer a tractable model system to study Obelisk molecular biology. With 15 exemplar Obelisk sequences (Figure 4), an “Obelisk blueprint” arises: a ∼730-1340 nt apparently circular RNA; with an extended “rod-like”, largely symmetrical predicted RNA secondary structure (Figure 4 - “jupiter” plots); an Oblin-1 homologue whose RNA sequence is largely self-complementary (which `ColabFold` predicts occupies a “globule-like” tertiary structure in 9 of 15 examples, Figure 4 - tertiary structures); and an occasionally present second, smaller protein (e.g. Oblin-2).

Many questions arise about the Obelisks. Does their transmission involve a separate, more complex, infectious agent (like HDV)? Do they primarily spread via virus-like particles, or cytoplasmically like viroids? Are Obelisks plasmid-like in that they can co-exist, and in some cases, contribute to host adaptability and fitness? Like viroids and HDV, do Obelisks replicate via rolling circle replication using a co-opted host RNA polymerase? What roles do the apparently circular Obelisk genome topology and the evidently conserved Obelisk genomic secondary structure play in the Obelisk lifecycle? Is Oblin-1 an RNA binding protein, and how does *domain-A* factor into its function? Does Oblin-2 act as a competitive inhibitor of host leucine zippers, as a multimerizing element, and/or can it interact with Oblin-1? How do Obelisks that lack Oblin-2 complement its function(s)? What role do the Obelisk-specific self-cleaving ribozymes play, and how do they interact with the Oblin proteins? How do Obelisks affect their host, and are they largely a deleterious or beneficial element to harbour? And what impact, if any, does harbouring an Obelisk have on ‘meta’-host physiology, is Obelisk positivity predictive of human health states?

Lastly, Obelisks do not closely resemble any existing mobile genetic elements, raising the question of their appropriate designation. Throughout this work, Obelisks have been referred to as ‘viroid-like’, drawing comparisons to viroids and HDV. However, viroids are in-part defined by their non-coding nature ^6,7^, and HDV-like elements are defined by homology to the large hepatitis Delta antigen (L-HDAg, and in the case of HDV, human tropism and a satellite relationship to Hepatitis B virus) ^8^. By virtue of their predicted coding capacity, which does not resemble L-HDAg, Obelisks are then neither strictly viroids, nor Delta-like elements. The predicted self-complementarity of the Oblin-1 further deviates from L-HDAg, likely imposing a set of unique evolutionary constraints (protein tertiary structure in addition to RNA secondary structure), that are not experienced by viroids and HDV-like elements. We therefore propose these proteins be referred to as “Oblins”. Viruses are already ill-defined, with sub-viral agents (such as viroids and HDV) being defined within the then more nebulous ‘perivirosphere’ ^45^, but part of ‘sub-virality’ is the implication of virus-like behaviour, either in transmission (*e.g.* via virions), in host impact (*e.g.* a pathology), or in replication (*e.g.* a co-opting viral replication machinery). Currently, it is not possible to assign transmission mode, host impact, or replication mode of Obelisks, suggesting that these elements might not even be ‘viral’ in nature and might more closely resemble “RNA plasmids”. As such, we propose that the term “Obelisk” be used to refer to these agents as they are distinct from other sub-viral satellites ^46^, viroids, and HDV.

## Methods

### VNom

`VNom` (pronounced *venom*, short for “Viroid Nominator”) was written to sequentially filter, in a homology-independent manner, for contigs with molecular features consistent with viroid-like biology from *de novo* assembled stranded RNA-seq data, namely: apparent circularity, and the co-occurrence of both positive- and negative-sense strands within a given sample (Supplementary Figure 2). As an input, `VNom` can take in any De Bruijn graph assembled contigs from stranded RNA-seq data; however, `VNom` is optimised to work on the output from `rnaSPAdes` ^47^. Initially, apparent circularity is inferred by identifying perfect k-mer repeats between the start and end of a contig: a previously exploited ^2,12,48^ sequence feature produced from circular De Bruijn graphs which are in turn produced from repetitive or circular transcripts during assembly. These apparently circular contigs are further de-concatenated into apparent unit-length, monomeric sub-sequences if a regular repetition of the identified k-mer is found, as is analogously done in ^14^. The resulting apparently circular contigs are then clustered with `circUCLUST` ^49^ and clusters containing at least one apparent sense and one antisense contig are kept (as inferred by k-mer counting). Any previously filtered out contigs that produce strong global alignments (`usearch -usearch_global`) ^50^ to these resulting sense-antisense clusters are then re-introduced where any clusters with now mutual contigs are merged. Local alignment (`usearch -usearch_local`) is then used to resolve and annotate any new multi-unit-length contigs into monomeric sub-sequences, and any sub-unit-length sequences into fragments. Finally, the resulting clusters are all “phased” to the same circular permutation using the multiple sequence aligner `MARS` ^51^. `VNom` is freely available at github.com/Zheludev/VNom.

### Initial Obelisk identification

Stranded RNA-seq data were fetched from the SRA ^52^ using `fasterq-dump` ^53^, adapter and quality filtered using `fastp` (`--average_qual=30 --n_base_limit=0 --cut_front --cut_tail`) ^54^, and *de novo* assembled with `rnaSPAdes` (default settings). Viroid-like sequences were identified using `VNom` (`-max 2000-CF_k 10 -CF_simple 0 -CF_tandem 1 -USG_vs_all 1`).

Obelisk RNA was initially identified in a longitudinal dataset of human stool stranded metatranscriptomics from the Integrative Human Microbiome Project (iHMP) ^20^. All paired-end RNA-seq datasets were downloaded (104 donors), trimmed, and assembled as described. Contigs were then grouped by donor ID and passed through `VNom`. The 2306 resulting `VNom`-nominated sense contigs were then queried manually for apparent lack of nucleotide, or protein-coding homology to the NCBI `nt/nr` (see later in this paragraph). Amongst these, we chose a sequence with striking predicted RNA secondary structure (high degree of basepairing, by eye, `RNAfold -p -d2 --noLP --circ`) ^55^. Obelisk RNAs were also manifest when `VNom` nominated contigs were passed through the following pipeline: the sense contigs were queried against a custom database (see Data Availability) of self-cleaving ribozymes (`CMscan`, default settings, keeping any, including likely spurious, hits) ^56^, these resulting 196 contigs were then assayed against the NCBI `nt` database (*11 Oct 2021*, `blastn`, default settings) ^21,57^, and contigs that yielded no hits, or whose best (by E-value) hits aligned to less than 40 % of contig’s length were kept. These resulting 20 contigs were then queried against the NCBI `nr` database (*8 Nov 2021*, `blastx`, default settings), similarly keeping sub-40 % alignment length best hits, yielding 11 contigs, of which 5 had a unit length of 1164 nt (one contig was 1166nt) - suggesting a common class of RNA. These were later defined as the Obelisk RNAs. Similarly, `blastn/p` filtering the 2306 sense contigs but without the `CMscan` step yielded 107 contigs, 8 of which were over 1000 nt in length, comprising the 6 Obelisk RNAs. Lastly, running `blastn` on all the iHMP contigs against the 6 Obelisk RNAs resulted in a final total of 15 unique Obelisk RNA sequences.

### Taxonomic classification

Taxa from length-filtered reads (fastp, as above with --length_required 75) were classified using Kraken2 (default settings) ^58^ against the Phanta ^59^ database, modified with non-redundant Obelisk-ɑ/β sequences using KrakenGrafter ^60^, followed by Bayesian re-estimation using Bracken (-r 75) ^61^, lastly taxon counts were combined using Bracken2OTU ^60^, summing any samples that came from the same donor on the same day (indicative of split sequencing lanes).

Obelisk-ɑ positive length-filtered read datasets, were assessed for sequence diversity relative to a fixed, arbitrarily chosen Obelisk-ɑ reference ^62^. Namely, single nucleotide polymorphisms (SNPs) and small structural variants were measured by aligning reads (`bwa-mem2`, default settings) ^63^ to the reference, followed by deduplication (`picard`, `MarkDuplicates`) ^64^, and detection `freebayes` (`--ploidy 1--pooled-discrete --pooled-continuous)` ^65^. `SAMtools` ^66^ and `bamaddrg` ^67^ were used throughout. Principal component analysis (PCA) on the resulting `vcf` file was computed using `SNPRelate` (`snpgdsPCA`)^68^, as described in ^69^, clusters were identified by `kmeans` (`centers = 5`) ^70^.

### Obelisk homologue detection in additional public data

Close Obelisk-ɑ homologues were identified in the Short Read Archive (SRA) ^52^ using `PebbleScout` (“Metagenomic” database, default settings) ^25^, a recently released tool that efficiently queries ∼3.2 million (*mid 2022*) raw sequencing data for exact 42 k-mer matches. 9 metatranscriptome BioProjects (comprising 34 short read datasets) were identified (`PBSscore` > 65) with close (∼1 % nucleotide divergence) matches to Obelisk-ɑ, of which 3 were part of iHMP or its predecessor ^71^, 5 were from other human stool studies ^72–76^, and 1 was from a fox gut autopsy ^77^. Using the `VNom` pipeline (see above), 21 datasets (from 7 BioProjects) yielded full length Obelisk-ɑ sequences, all from human hGMB studies (Table 3).

Finding Obelisk-ɑ homologues in studies separate from the iHMP lent support to these RNA elements being legitimate biological entities. Further, one Obelisk-ɑ homologue was found in a study from our own institution ^76^, suggesting that Obelisk-like RNAs could be locally present. Emboldened by this, we solicited hGMB stranded RNA-seq data from the local academic community and identified closely related Obelisk-ɑ homologues in a dataset that at the time had not been uploaded onto the SRA (now available at `PRJNA940499`: donors `D01` - both Obelisks -ɑ and -β; and `D10` - just Obelisk-ɑ) ^78^. Further, within this dataset we identified a diverged Obelisk-like sequence with similar: length (1182 nt), lack of apparent homology to reference databases, predicted obelisk-like secondary structure, and two ORFs but with low homology to Obelisk-ɑ. In comparison to Obelisk-ɑ, this new “Obelisk-β” had a 41.30 % nucleotide sequence identity, and 23.42/38.29 % and 18.75/31.25 % on the amino acid level identities/similarities for ORFs 1 and 2, respectively (see below, Supplementary Figure 3a/c).

Owing to their apparent sequence novelty, the Obelisk-ɑ/β Oblin-1 and -2 protein sequences were next used as hallmark sequences specific to Obelisk-like RNAs - analogous to the use of RNA-dependant RNA polymerase (RdRP) hallmark sequences in RNA viral discovery ^1–5^. To identify divergent Obelisk-like elements, we searched the `RNA Deep Virome Assemblage` (`RDVA`, `v0.2`) ^13,26^, a collection of 58,557 assemblies of ∼12.5 trillion contigs, with `diamond` (`--very-sensitive`) ^79^ using Obelisk-ɑ/β Oblin -1 and -2 protein sequences deduplicated at 90 % sequence identity (`UCLUST`, default settings) as queries ^50^. This resulted in 38,545 sub-5000 nt hits which when de-replicated, circularly clustered (`circUCLUST`) into 29,859 and 19,808 clusters at 90 % and 75 % nucleotide sequence identity, respectively (see Data Availability). A conservative database of 7,202 Obelisks was built by keeping assemblies with a `CircleFinder` (`VNom` defaults) implied circularity, with each genome “phased” to 50 nt from the start codon of its largest predicted ORF (`prodigal`, `-p meta`). This database was clustered (`circUCLUST`) into 1,744 80 % identity clusters which were then sub-clustered at 95 % identity (Supplementary Table 1). The assemblies were then named based on these nested clusterings. A naming convention is proposed with the following pattern *“Obelisk_X_Y_Z”* where “X” refers to the 80 % cluster ordinate, “Y” to the 95 % cluster ordinate, and “Z” as a unique identifier within the 95 % cluster. The first 15 80 % ordinates are defined as the Obelisks depicted in Figure 4, the next 10 80 % ordinates are defined as the remaining letters in the Greek alphabet (*omicron* through *omega*). As such, the centroid Obelisk-ɑ sequence that is also the centroid of the first 95 % sub-type is defined as “*Obelisk_000001_000001_000001”*.

### Serratus

Extending from the `RDVA` search, a larger breadth of public datasets (5,470,176 runs) was next assessed for diverged Obelisk-like sequence presence. Profile hidden Markov models (pHMMs) of ORFs 1 and 2 were derived from the `RDVA` hits (see below) and used as queries in the `Serratus` architecture ^2^, an optimised, cloud-based pipeline for efficiently identifying sequencing reads that align to pHMMs. By looking for pHMM matches, `Serratus` is able to find more distantly related Obelisk-like sequences where k-mer match searches (*e.g.* `PebbleScout`) would fail, but at a considerable computational expense. Datasets were defined as a `Serratus` hit if at least one read aligned (E-value <1×10^-4^) to either Oblin-1 or Oblin-2. Of the resulting 949,810 non-redundant SRA hits, 215,398 datasets were selected by filtering with a virus-presence score (≥25, explained in github.com/ababaian/serratus/wiki/.summary-Reports) which attempts to predict ORF *de novo* assembly success, ultimately yielding 1,499 datasets containing both Oblin-1 and Oblin-2, 3,006 containing only Oblin-1, and 213,891 containing only Oblin-2. Per hit SRA, high confidence ORF mapping reads were then *de novo* assembled using `rnaSPAdes` (default settings) yielding Obelisk “micro-assemblies”. This `Serratus` run was conducted along with other pHMM queries, meaning that *de novo* assembly happened in aggregate with all other hits, as such, `diamond` (`--very-sensitive`) was used to extract Oblin-1/-2 micro-assembly protein sequences.

### Protein homology bioinformatics

To probe the deep sequence diversity of Oblins 1 and 2, corresponding single domain profile hidden Markov models (pHMMs) were individually constructed from the `RDVA` hits using an iterative approach: A multiple sequence alignment (MSA) from the initial `PebbleScout` set was computed using `Muscle5` (default settings) ^80^, from which an initial pHMM was computed using `HMMbuild` (default settings) ^81^. Each genome in the `RDVA` non-redundant 90 % sequence identity cluster centroid set was doubled in length using `SeqDoubler` ^60^ and ORFs were predicted using `Prodigal` (`-p meta`) ^82^. ORFs with predicted N- or C- terminal truncation were omitted and a non-redundant set was kept (`usearch -fastx_uniques`) ^50^. This ORF database was queried against (`HMMsearch,` default settings) the initial pHMM and hits with global E-values lower than 1×10^-15^ for Oblin-1 or 1×10^-8^ for Oblin-2 were kept. `HMMalign` (`--trim`) and `MSACleaner` (`-ref` from the `PebbleScout` set and `-fxn 0.01`) ^60^ were used recursively (until no new sequences were omitted) to filter the constituent MSA sequences to omit sequences that contributed large indels relative to the initial pHMM. A new pHMM was computed and the `HMMsearch` (on the remaining ORFs), `HMMalign` (without `--trim`), and `MSACleaner` steps were repeated once. This resulting MSA was filtered by sequence length `FASTACleanUp` (`-lower` 150 for Oblin-1, `-lower` 40 for Oblin-2) ^60^ and a final pHMM was computed. `msaconverter` ^83^ was used throughout. There were no overlapping sequences between the resulting Oblin-1 and -2 pHMMs.

A contiguous alignment block of 18 amino acids was noticed in the resulting Oblin-1 pHMM (Obelisk-ɑ: 152-RRRGYKDHGSRRFPHEVH-169) and was selected as a marker sequence, terming it *domain-A*. Because the `Serratus` Oblin-1 micro-assemblies may include some that are not full-length (*wrt* Oblin-1), further aggregation from the `Serratus` data utilised a search for similarity to *domain-A*. To incorporate the `Serratus` results, an initial 503 sequence *domain-A* alignment was extracted from the `RDVA` pHMM (and later used with `K-mer Rabbit`, below) and a new pHMM was constructed (`HMMbuild`, default settings). A length sorted (`seqkit sort -l -r`), non-redundant (`usearch -fastx_uniques`) set of `Serratus` Oblin-1 micro-assemblies was then iteratively queried with an ever-rebuilt *domain-A* pHMM: keeping `HMMsearch` (default settings) hits with E-values lower than 1×10^-4^, intermediate MSAs were re-built (`HMMalign --trim`) relative to the previous iteration and sequences with at least 8 amino acids (`seqkit seq -g -m` 8) were kept, next, the resulting sequences were re-aligned to the current pHMM and a new pHMM was built, lastly, all <1×10^-4^ E-value hits were omitted and a new iteration was started. A finalised `Serratus`-inclusive *domain-A* pHMM was constructed with 30,686 sequences after 12 cycles. This process was repeated for two other less well-conserved domains, *domain-B* (Obelisk-ɑ: 96-CLTSKSGMLNFLEDTTLY-113), and *domain-C* (Obelisk-ɑ: 53-RSKKDLLALAIISWWLEE-70), with 5076 and 5103 resulting sequences, respectively. *Domains -B/-C* were not studied further in this work.

### Protein tertiary structure prediction

For initial, monomeric tertiary structure prediction, `RDVA` pHMM MSAs were re-aligned (`Muscle5`, default settings) relative to ORFs-1/2 from Obelisk-ɑ and used with `ColabFold` (`v1.5.2-patch`) ^32^ implementation of `AlphaFold2` (default settings, no `amber`, no `dropout`) ^31^. The `HHblits` suite was used to convert between `fasta` and `a3m` MSA formats ^84^. Tertiary structure homology was assessed using the `Phyre2` (default settings) ^85^, `Dali` (`PDB Search`) ^86^, `FoldSeek` (all databases, `3Di/AA` and `TM-align` scoring) ^87^, and the `Clustered AlphaFold Database` ^88^ webservers (see Data Availability). For all other tertiary structure predictions, `ColabFold` was used with `mmseqs2 uniref env` for MSA generation. For 9 in 15 predictions, including Obelisks -ɑ, -β, and -*S.s*, this yielded qualitatively similar “globule” predictions (Figure 4 - tertiary fold predictions). An equivalent 73 sequence MSA was constructed for Oblin-1 homologues from ribozyme-baring Obelisks (see RNA homology bioinformatics) by first filtering any `Prodigal`-predicted proteins for length (`seqkit seq -m 200 -M 250`), aligning the resulting sequences (`Muscle5`), and manually removing any sequences that appeared to disrupt the MSA. `ColabFold v1.5.3` was used for ribozyme-baring Oblin-1 protein tertiary fold predictions and Obelisk-nu.

### Protein conservation and phylogenetics

Oblin-1/-2 conservation analysis was conducted on Obelisk-ɑ-relative `a3m` alignments against the `BLOSUM62` substitution matrix ^89^ using `msaConservationScore` (`gapVsGap` = 0) ^90^ and the `Biostrings` package ^91^. The Oblin-2 sequence logo was constructed using `ggseqlogo` ^92^, and a consensus sequence was generated with `msaConsensusSequence` (`upperlower`, `thresh` = 20,0).

Owing to the micro-assembly used in the `Serratus` search, phylogenetic analysis was limited to the highly conserved *domain-A*. To ensure a *domain-A* phylogenetic tree encompassed the observed sequence diversity from ribozyme-baring Obelisks, the underlying multiple sequence alignment (MSA) construction started with an iterative pHMM construction approach similar to method used to build the initial Oblin-1 pHMM. First Oblin-1 homologues from ribozyme-baring Obelisks (see RNA homology bioinformatics) were queried (`HMMsearch --max`, E-value ≤ 1×10^-8^) against the initial Oblin-1 pHMM, yielding only sequences homologous to *domain-A*. These sequences were re-aligned (`Muscle5`) and an initial ribozyme-associated *domain-A* pHMM was built. This ribozyme-associated pHMM was then iteratively built upon with successive rounds of similarity searches (`HMMsearch --max`, E-value ≤ 1×10^-8^) against the `RDVA`’s ribozyme-baring Obelisk’s predicted proteins followed by re-alignment with `Muscle5`. Once no new sequences were found, the cycle was continued at an E-value threshold of 1×10^-5^. This resulting ribozyme-associated MSA was then re-aligned to the initial Oblin-1 MSA (`HMMalign`, default settings) and the alignment column corresponding to *domain-A* was manually excised, and re-aligned (`Muscle5`). The entirety of the full-length predicted proteins from the `RDVA` were then similarly iteratively queried but at a E-value threshold of 1×10^-4^, and without an intermediate `Muscle5` step. The converged alignment was then re-aligned with `Muscle5` (`Super5`) and similarly iteratively queried against the `Serratus` micro-assemblies, keeping the best hit per micro-assembly until convergence. The resulting 46,884 total *domain-A* sequences were finally re-aligned with `Muscle5` (`Super5`). This MSA was then deduplicated, and optimised using `CIAlign` ^93^ to remove insertions (minimum size 1, minimum 0.05 %), to crop divergent sequences (minimum identity proportion 0.01, minimum non-gap proportion 0.5, buffer size 4), and to remove any resulting sequences shorter than or equal to 16 aa. A final round of deduplication yielded a 3265 non-redundant sequence *domain-A* no-gap alignment of 17 aa (Supplementary Table 2).

A maximum likelihood phylogenetic tree was then constructed from this 17 aa alignment using `iqtree` ^94^. The `LG+G4` substitution model (`testnewonly`) was selected (`ModelFinder` ^95^) based on a consensus between the Akaike and Bayesian Information Criteria. Tree construction was run with 33,000 UFBoot bootstraps ^96^, Nearest Neighbour Interchange optimization, and 33,000 SH-like approximate likelihood ratio tests (`-B 33000 -bnni -alrt 33000`). The resulting tree was plotted using `iTOL` ^97^.

### ScanRabbit

For rapidly searching smaller, locally-held datasets for novel Obelisk homologues, we developed a second tool, `ScanRabbit`, which focuses on a short segment of any multiple sequence alignment. `ScanRabbit` was run using the position-specific-scoring matrix (PSSM) based on the multiple sequence alignment used to build the Oblin-1 profile hidden Markov model (see above) from the `RDVA` hits corresponding to *Domain-A*. `ScanRabbit` accelerates searches on local hardware through direct bitwise conversion of the PSSM to a local bitwise scoring that can be applied to the raw binary representation of RNA-seq reads, and a just-in-time compiler `PyPy` ^98^. `ScanRabbit` is available on GitHub at github.com/FireLabSoftware/ScanRabbit.

### Obelisk spacer analysis

The presence of Obelisks in known prokaryotic CRISPR spacer arrays was assessed using a conservative k-mer matching approach. Namely, the `RDVA` Obelisk dataset was queried against predicted CRISPR spacers in the Joint Genome Institute (JGI) `IMG/M` spacer database (*May 2023*) ^36^. To estimate a lower length bound on matching noise, a parallel analysis was conducted on “reversed” (*not* reverse complemented) Obelisk sequences. Initially, `RDVA` Obelisk sequences were searched against the `IMG/M` spacer database using `blastn` (default settings), only keeping perfect matches with no gaps or mismatches (k-mers) - the longest k-mer match between a given spacer/Obelisk pairing was kept. Next, all kept spacers containing any 12-mer match to common Illumina sequencing adaptors were omitted using `KmerCatcher` (default settings) ^60^. For each remaining spacer, the information content was estimated ^99^ by comparing how efficiently the compression algorithm `zip` (`-9`) ^100^ could “deflate” a given spacer - a larger length normalised deflation indicates a less complex spacer sequence that is less likely to be unambiguously mapped to a specific (Obelisk) sequence. The repetitive content of each spacer was also assessed using `etandem` (`-minrepeat` 4, `-maxrepeat` 15, `-threshold` 2) ^101^. Spacers with a length normalised deflation less than 1.0 percent per nucleotide were kept (137,667 forward, 118,411 reverse), these spacers also qualitatively had a low `etandem` score though this metric was not used for filtration (Supplementary Figure 8a). Next, only the 23 forward spacers longer than the maximum length of the reverse spacers (25 nt) were kept as any mappings below this threshold would be indistinguishable from noise (reverse-mapping, Supplementary Figure 8b). Lastly, the corresponding Obelisks mapping to these spacers were minimum length filtered to 1000 nt (`seqkit seq -m` 1000), resulting in two contigs. Only one of these contigs gave `blastn` (default settings, `NCBI` webserver, *August 2023*) a largely (∼95 %) unknown sequence with a singular ∼45 nt sequence mostly showing up in high G+C Gram-positive bacteria and cyanobacteria (consistent with a CRISPR spacer array, see Data Availability). This largely unknown 1096 nt contig was found to encode (`prodigal -p meta`) homologues of both Oblin-1 and Oblin-2 (`HMMsearch`, default settings, against the `RDVA` pHMMs), and is predicted to fold (see below) into an obelisk-like RNA secondary structure (Supplementary Figure 8c) - features consistent with being an Obelisk which we term Obelisk-“gamma” (Obelisk-ɣ). Two spacers were found to map to Obelisk-ɣ, both from the same *Bombella mellum* genome (`RefSeq GCF_014048465.1`) ^102^ - these spacers (which differ by one extra nucleotide) were found at the same putative CRISPR locus but predicted in the `IMG/M` database with two different tools (`PILER-CR` and `CRT`) ^103,104^, as such, this is likely one spacer. Obelisk-ɣ’s predicted secondary structure is not as “rod-like” as other Obelisks (Figure 4 - “jupiter” plots), with the spacer mapping to the “frayed” end; additionally, the spacer mapping position coincides with the locus identified by `blastn`; and lastly, `CircleFinder` (`VNom` default settings) did not identify a start-end k-mer repeat indicative of a circular genome. The Obelisk-ɣ Oblin-1 was also not predicted (see above) to fold into the characteristic “globule” fold (Figure 4 - tertiary fold predictions), though the discriminatory power of this is unclear and so ignored. These features suggest that the Obelisk-ɣ genome might be mis-assembled, with the putative spacer mapping sequence arising from a chimeric assembly. As such, this conservative approach to CRISPR spacer mapping was not able to unambiguously identify any Obelisk relationships to CRISPR spacer arrays as we currently recognise them.

### Identity and similarity measurements

Unless otherwise stated, all nucleotide identity, and protein identity and similarity measurements were computed by first building a pairwise alignments Muscle5 (default settings) of “phased” genomes (as below) followed by calculation with Ident and Sim (default settings) ^105^.

### RNA homology bioinformatics

Figure 1b, Figure 4, Supplementary Figure 3b, and Supplementary Figure 6a RNA secondary structures were predicted using `RNAalifold` (`-p`, `-r`, `-d2`, `--noLP`, `--circ`) ^107^ on the non-redundant (`usearch-fastx_uniques`), 1164 nt long, `PebbleScout` set of the above “phased” Obelisk-ɑ sequences, split by genome polarity, using a `Muscle5` (default settings) derived MSA. Supplementary Figures 1a and 2c-d secondary structures were predicted on singular genomes using `RNAfold` (`-p -r -d2 --noLP --circ`). RNA secondary structures were illustrated using `circlize` ^108^ for “jupiter” plots, and `R2R` ^109^ for “skeleton” diagrams. Conserved RNA element (*e.g.* ribozymes) coordinates in Supplementary Figure 1 were identified using `CMscan` (`--rfam --cut_ga`) against the `Rfam 14.6` database ^110^.

23 Obelisk-encoded hammerhead type-III ribozyme homologous sequences were initially identified (`CMsearch`) using the `RF00008` reference covariance model against the 90 % identity-clustered (`circUCLUST`), sequence-doubled (`SeqDoubler`) `RDVA` dataset, using stringent cutoffs for confident (E-value ≤ 1×10^-5^), full-length (`--notrunc`) hits, keeping only the best hit per Obelisk genome. An Obelisk-specific, “Obelisk-variant hammerhead type-III” (ObV-HHR3) covariance model (CM) was constructed using an iterative approach: an initial CM was constructed using the 23 hit sequences by aligning them against `RF00008` (`CMalign`, default settings), optimising the alignment using `CaCoFold` ^111^ (`R-scape`: `-s`, `--cacofold`, `--rna`), and finally building (`CMbuild`, default settings), and calibrating (`CMcalibrate`, default settings) the CM. Using this initial CM as a starting point, the sequence-doubled `RDVA` dataset was iteratively passed through the `CMsearch`, `CMalign`, `CaCoFold`, `CMbuild`, and `CMcalibrate` pipeline, each time only keeping the best, non-truncated, E-value ≤ 1×10^-5^ hits (one hit per Obelisk genome) and additively appending them to the CM, subtracting the hits from the `RDVA` set as they were found, until no new hits were found. Ultimately, a 178 sequence ObV-HHR3 was constructed with 15 significantly covarying positions identified (Supplementary Figure 6b). When re-querying (`CMsearch`, `--no-trunc`) the full `RDVA` dataset with this finalised CM at an E-value ≤ 1×10^-4^, 339 Obelisk genomes were identified. The ObV-HHR3 column in Supplementary Table 1 was annotated with `CMsearch`, `--no-trunc`, ≤ 1×10^-5^ on sequence-doubled genomes.

### Streptococcus sanguinis bioinformatics

In an attempt to identify Obelisk-like elements that had been serendipitously sequenced in isolation with their putative cellular host(s), Oblin-1 positive filtered `Serratus` hits were screened for potentially low biodiversity experimental designs such as defined co-culture, single-cell RNA-seq, and isolate culture. As such, isolate RNA-seq experiments of *Streptococcus sanguinis* (strain SK36, a commensal of the human oral microbiome) stood out (Table 4). Upon further investigation (using `CircleFinder` from `VNom`), a 1137 nt, obelisk-shaped RNA coding only for Oblin-1 was identified. This so-called “Obelisk-*S.s*” exhibited 40.65 % and 35.47 % nucleotide sequence identity with Obelisk-ɑ and Obelisk-β, respectively, and 19.92/33.47 % and 21.05/32.71 % Oblin-1 amino acid identity and similarity to Obelisk-ɑ and Obelisk-β, respectively (see above, Supplementary Figure 3a/d). Additionally, Obelisk-*S.s* was further found in human oral microbiome samples (Table 5, Supplementary Figure 9), and by comparing isolate cultures from different growth media, *S. sanguinis* was determined to be the likely cellular host as opposed to Obelisk-*S.s* being a contamination from complex media.

### Surveying for Obelisks in human data

The prevalence of Obelisks in five human microbiome datasets (three gastric, hGMB, and two oral, hOMB, Table 5 and Table 6) was (re-)evaluated after both Obelisks -ɑ, -β, and -*S.s* were identified, and the `RDVA` pHMMs were constructed. For human gut metatranscriptome data, the 104 iHMP donors ^20^, and the 10 “ZF” donors from the dataset where Obelisk-β was found ^79^ were reanalysed; additionally, 326 new donor samples from an irritable bowel syndrome study ^112^ were queried, for a total of 440 hGMB donors analysed. For human oral metatranscriptome data, 22 (50/50 healthy/case) donors from a Dutch cohort studying periodontitis ^37^, and 10 healthy donors from a oral extracellular vesicles study ^113^ were queried for a total of 32 hOMB donors analysed. To identify more diverged Obelisk elements, a pHMM mapping approach was taken - similarly to `Serratus`. Namely, each dataset’s trimmed reads (as before) were translated in all six frames (`seqkit -f` 6`-F`) and assessed for Oblin-1 homology using `HMMsearch` (default settings) against the `RDVA` Oblin-1 pHMM. Donors with greater than or equal to 10 translated reads (averaging over per-donor replicates, time points, or sampling locations if present) mapping with an E-value less than or equal to 1×10^-5^ were counted as true Oblin-1 hits. Additionally, these trimmed reads were assessed for Obelisk -ɑ, -β, and -*S.s* presence using a modified `Phanta Kraken2` and `Bracken` database constructed as before incorporating all non-redundant Obelisk -ɑ, -β, and -*S.s* sequences (only the previous Obelisk-ɑ positive iHMP datasets were re-assessed in this way). Across these five datasets, 21 donors were identified as positive for Obelisk homologues (>10 `HMMsearch` hits) but negative for Obelisks -ɑ, -β, or -*S.s* (<10 `Kraken2` hits), additionally, 25 donors were identified as positive for Obelisks -ɑ, -β, or -*S.s* (>10 `Kraken2` hits, Supplementary Figure 9). The presence of pHMM-mapping reads in the absence of k-mer reads suggested the existence of new Obelisks, as such, these 21 donors’ datasets were assessed for new Obelisks. Briefly, these donor’s trimmed reads were assembled as before, keeping any contigs with Oblin-1 homology (`HMMsearch`, `--max`, E-value ≤1×10^-5^), and then selecting for apparently circular contigs with `CircleFinder` (default `VNom` settings). These selected contigs were next assessed for Oblin-2 coding capacity (`prodigal -p meta`, followed by `HMMsearch` and `blastn` versus the Oblin-2 `RDVA` pHMM, and Obelisk-ɑ Oblin-2 sequence and consensus, respectively E-value ≤1×10^-4^), and obelisk-like secondary structure as before. Clustering all resulting and previously identified contigs (`circUCLUST -id` 0.8), 11 new full-length Obelisks were identified, which we named “delta” through “xi” (“*Obelisk_000005*” to “*Obelisk_000015*” in Supplementary Table 1, Figure 4). “Delta,” “epsilon,” “zeta,” and “eta” were found in the hGMB datasets and all remaining Obelisks were found in the Dutch hOMB dataset - indicating a human sampling site specificity to Obelisk species. Of these 11 Obelisks, eight apparently only code for an Oblin-1 homologue, Obelisk-“kappa” codes for an Oblin-2 homologue, and Obelisks -“lambda” and -“mu” code for a second ORF similar in size to Oblin-2 but with no obvious homology (which we term the “2ndORF” as more study is needed to determine if this is actually a *bona fide* new ORF). Four of these new Obelisks’ (“epsilon,” “kappa,” “mu,” and “xi”) Oblin-1 sequences were not predicted to fold (as before) into the otherwise characteristic “globule” tertiary structure (Figure 4 - tertiary fold predictions). These new Obelisks span between 733 nt (Obelisk- “iota”) to 1372 nt (Obelisk- “kappa”). Considering these new Obelisk sequences, as well as donors which did not yield full-length Obelisk candidates, Obelisks appear to occur in 9.5 % of the human donors assayed (6.6 % of hGMB samples, and 53 % of hOMB samples) and describe a wider breadth of characteristics that Obelisks seem to be able to possess (length and coding capacity).

**Table 6.** Supplementary Figure 9 donor aliases. Human donor aliases used for brevity in this study (Supplementary Figure 9) and their corresponding nomenclature from their original studies. Note, datasets from *Maghini and Dvorak et al. 2023* ^79^ and *Tong et al. 2023* ^113^ were used without re-assigning new aliases.

| study | alias to study nomenclature key |
| --- | --- |
| Lloyd-Price *et al.* 2019 <sup>20</sup> | A:2077, B:6038, C:2068, D:5001, E:2071, F:2025, G:3034, H:4016, I:4022 |
| Jacobs *et al.* 2023 <sup>112</sup> | A:A6405, B:A6673, C:1634K, D:A6089, E:1596C, F:A7010, G:A6942, H:A6889, I:A6419, J:A6797, K:A6083, L:A6974, M:A6131, N:A6739, O:A6810, P:A6745, Q:1558C |
| Belstrøm and Constancias *et al.* 2021 <sup>37</sup> | A:K2, B:K3, C:MP8, D:MP10, E:K11, F:K10, G:MP11, H:K9, I:MP1, J:K6, K:MP4, L:MP2, M:MP9, N:K7, O:K5 |

## Acknowledgements

This work is dedicated to the memory of mentor and friend, Paul Berg.

This work was funded by: Stanford Graduate Fellowship (INZ); University of Valencia Margarita Salas Fellowship MS21-067 (MJLG); Generalitat Valenciana Grant PROMETEO CIPROM/2022/21 (MDLP); Ministry of Economics and Competitivity of Spain-FEDER grant PID2020-116008GB-I00 (MDLP); Canadian Institutes of Health Project Grant PTJ-496709 (AB); Computing resources were provided by the University of British Columbia Community Health and Wellbeing Cloud Innovation Centre, powered by AWS (AB); NIH Grant R01AI148623 (National Institute of Allergy and Infectious Diseases, NIAID) (ASB); NIH Grant R01AI143757 (NIAID) (ASB); Convergence Grant 3.1416 (Stand up 2 Cancer) (ASB); Paul Allen Distinguished Investigator Award (ASB); and NIH Grant R35GM130366 (National Institute of General Medical Sciences) (AZF). We are grateful to the greater biology community for open data sharing. Some figures were arranged in BioRender.

## Data availability

Code and tabular summaries are available at the Stanford Digital Repository (purl.stanford.edu/wb363nt3637).

## Supplementary Figures

**Supplementary Figure 1.**
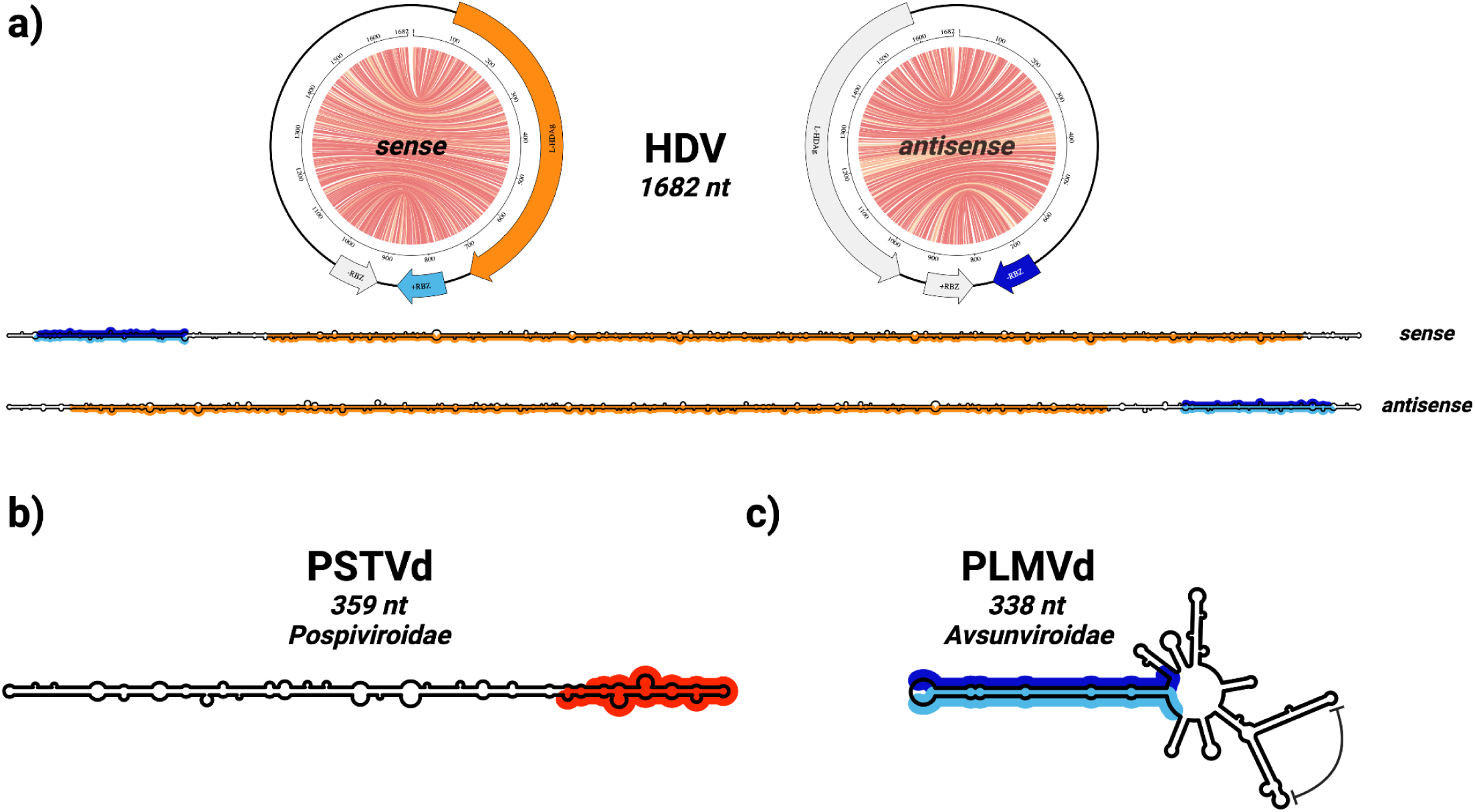
Background on viroid and HDV families: Hepatitis delta virus, *Pospiviroidae*, and *Avsunviroidae* form a class of highly structured, circular sub-viral RNAs. **a**) the Hepatitis delta virus (HDV) genome (`NC_001653.2`) ^114^ is predicted to fold into a rod-shaped RNA secondary structure in both sense, and antisense - depicted here as both “jupiter” plots where chords represent predicted basepairs (coloured by basepair probability from 0, grey, to 1, red) with features greyed out in antisense, and “skeleton” diagrams. Large hepatitis delta antigen (L-HDAg, orange), and hepatitis delta ribozymes (RBZ, Rfam: `RF00094`, antisense: dark blue, sense: light blue) indicated. **b**) Potato spindle tuber viroid (PSTVd) of the family *Pospiviroidae* folds ^115^ into a rod-like RNA secondary structure similar to HDV but encodes no ORFs, though does possess a conserved Pospiviroid RY motif (Rfam: `RF00362`, red). **c**) Peach latent mosaic viroid (PLMVd) folds ^116^ into a highly basepaired, but “branched” RNA secondary structure as is characteristic of the *Avsunviroidae* family. Type III hammerhead ribozymes (Rfam: RF00008, antisense: dark blue, sense: light blue) and “P8” pseudoknot (curved flat-headed arrow) illustrated.

**Supplementary Figure 2.**
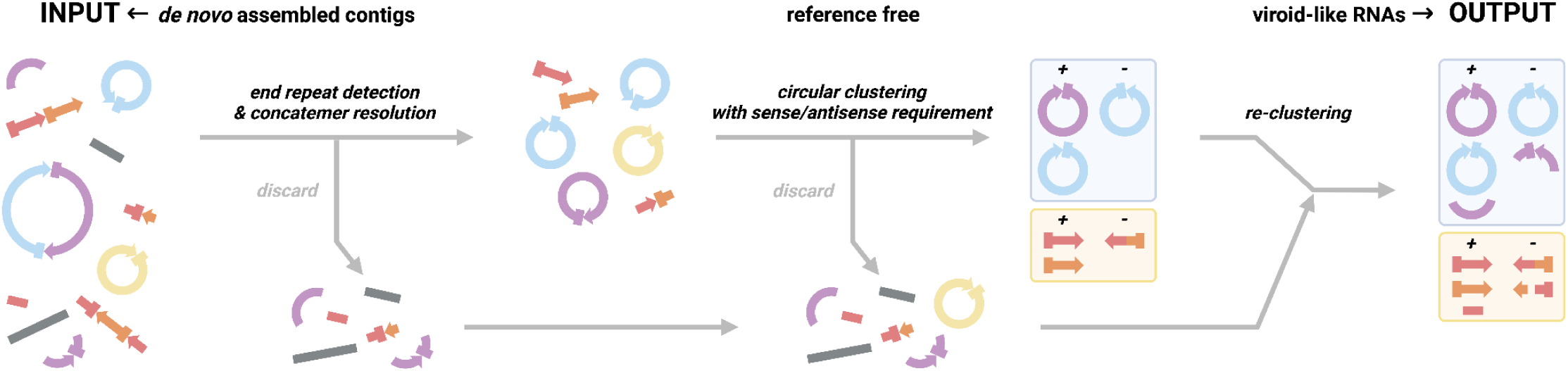
VNom sequentially filters contigs to enrich for RNAs with viroid-like properties. `VNom` (short for “Viroid Nominator”, pronounced *venom*) attempts to enrich for RNAs that are apparently circular and are present in the dataset in both polarities (a hallmark of RNA replication). To do this, `VNom` takes in *de novo* De Bruijn graph assembled contigs (from stranded RNA-seq data) and filters for potentially circular contigs by looking for perfect k-mer matches at the ends of each contig. Further, `VNom` also attempts to resolve concatemeric contigs by looking for regular repetition of such identified k-mers. These potentially circular contigs are then clustered based on sequence similarity using a circularly-permuting clustering algorithm. These resulting clusters are then kept if at least one contig of each polarity is identified by k-mer counting. Finally, these filtered clusters are compared against all of the previously discarded contigs to identify any remaining cluster members. While these filters should enrich for viroid-like RNAs, highly repetitive sequences also satisfy these requirements and so are often also enriched. `VNom` was found to work adequately well on deeply sequenced viroid-positive plant RNA-seq datasets (*e.g.* `SRR11060618`, `SRR11060619`, `SRR11060620`, and `SRR16133646`), especially when assemblies from the same bioProject were grouped together.

**Supplementary Figure 3.**
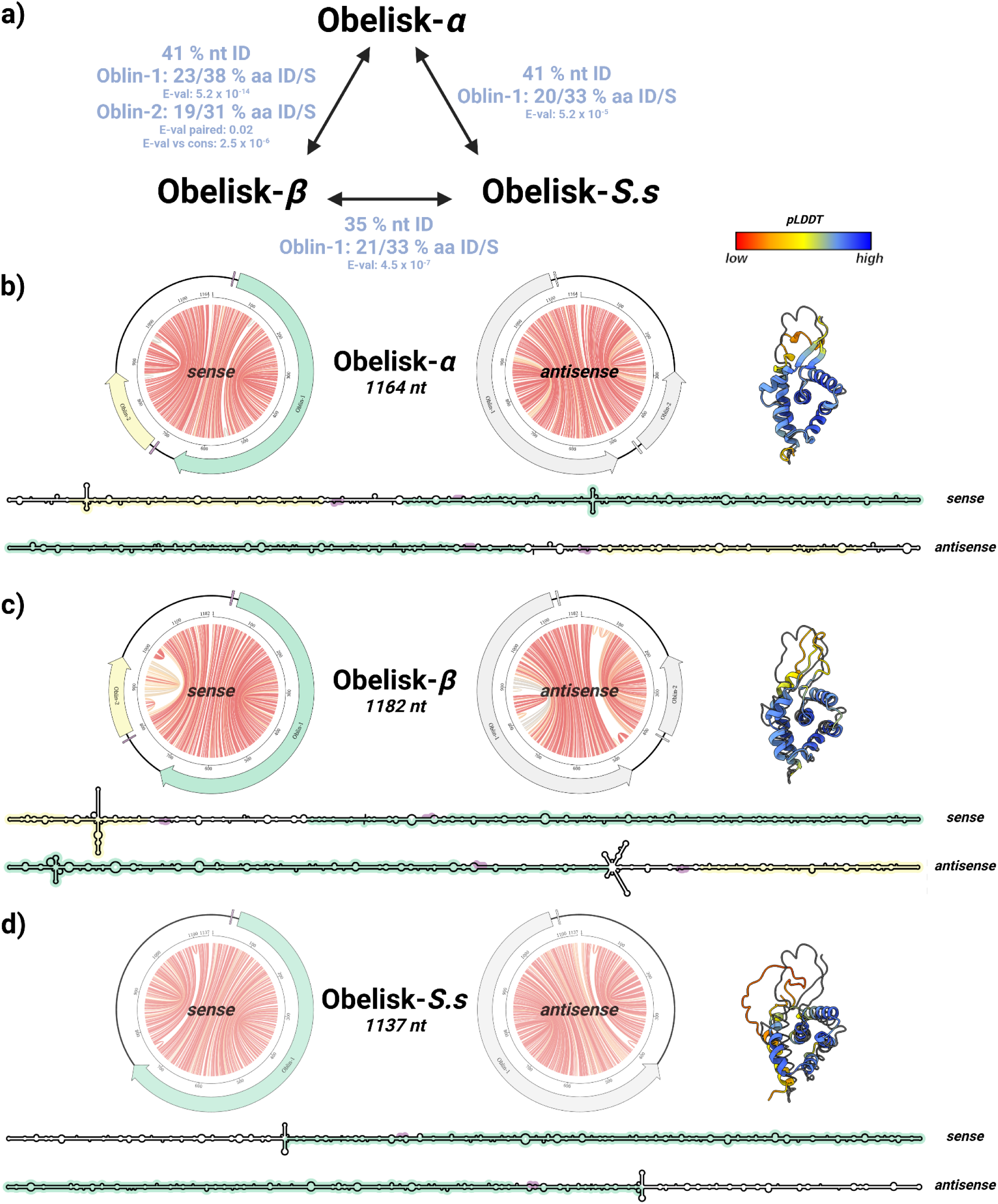
Obelisks -*alpha*, -*beta*, and -*S. sanguinis* appear to belong to the same, diverse family. **a**) nucleotide (nt) and amino acid (aa) -level pairwise sequence identities (ID) and similarities (S) between Obelisks- ɑ, β, and *S.s*. For Oblin protein sequences, mean pairwise `blastp` E-values are shown. Note, for Oblin-2 the pairwise `BLASTp` E-value relative to the Oblin-2 consensus (see methods) is also shown, indicating a distant, but evident homology between the ɑ and β Oblin-2s. **b-d**) These Obelisks are similar in lengths; 1164, 1182, and 1137 nt, respectively, and share globally similar obelisk-like predicted RNA secondary structures in both their sense and antisense - depicted here as both “jupiter” plots where chords represent predicted basepairs (coloured by basepair probability from 0, grey, to 1, red) with features greyed out in antisense, and “skeleton” diagrams. Likewise, the genomic synteny of predicted open reading frames (ORFs, preceded by predicted Shine-Delgarno sequences, purple) appear to be shared, with Oblin-1 (green) consistently being present on one half of the predicted RNA secondary structure, and Oblin-2 (yellow), when present, following shortly after Oblin-1. `ColabFold` predictions of Oblin-1 tertiary “globule” structures built with *ad hoc* multiple sequence alignment (MSA) construction (coloured cartoons) superimposed over the `RDVA`-derived MSA prediction for Obelisk-ɑ (black line, Figure 2a, see methods) indicating a conserved tertiary structure. Prediction confidence (pLDDT) shown as a colour bar (low confidence: 0, red; high confidence: 100, blue).

**Supplementary Figure 4.**
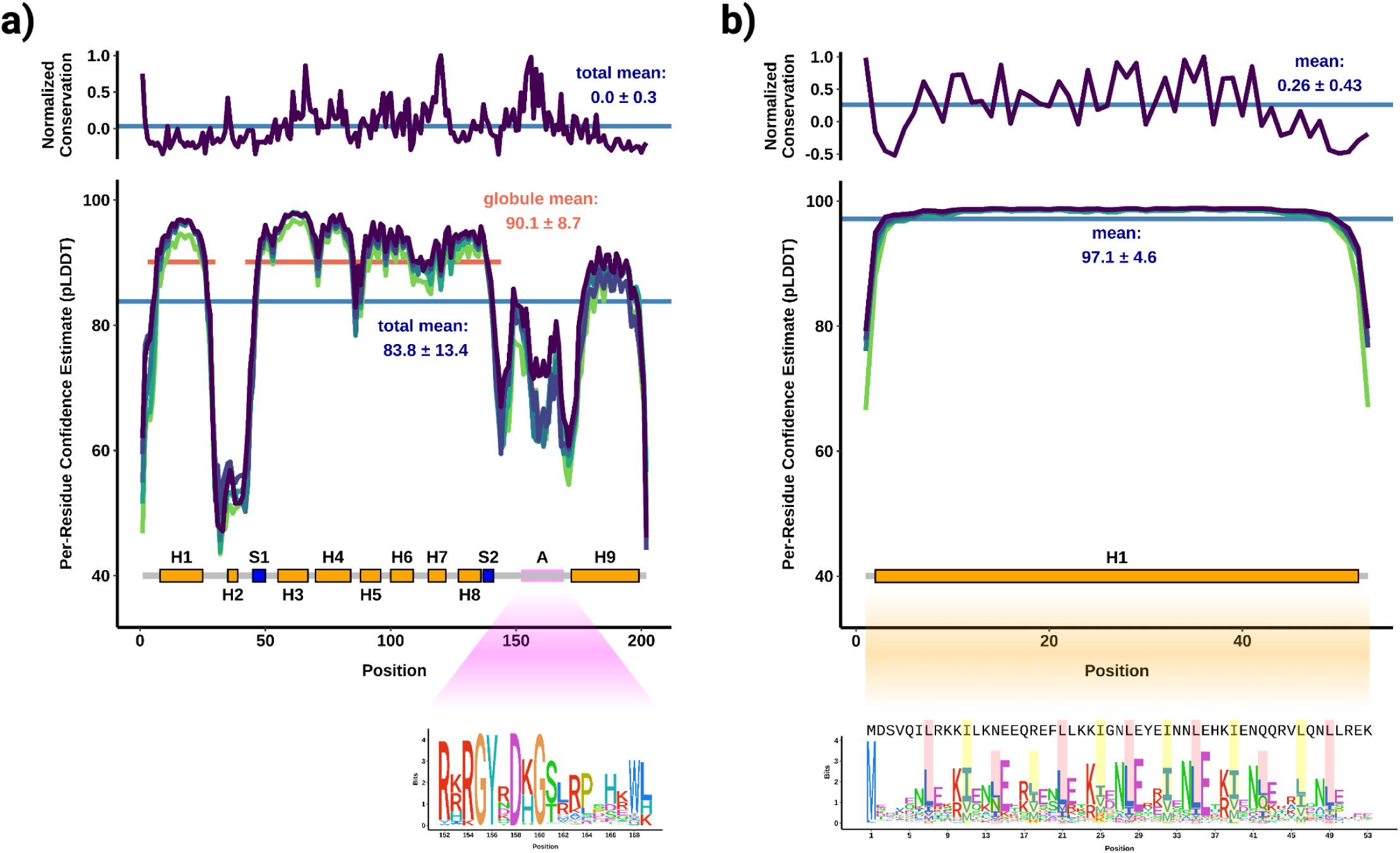
Oblins are diverse and generate robust protein fold predictions. **a**) normalised conservation (top, above zero = more conserved, see methods) of Obelisk open reading frame 1 (Oblin-1) relative to Obelisk-ɑ indicates that Oblin-1 is largely poorly conserved (mean per-residue confidence estimate, μ-pLDDT ± standard deviation of 0.0 ± 0.3) but has three regions of conservation, around the C-termini of alpha helices 3 and 7, and *domain-A* (see sequence logo callout, bottom). Oblin-1 tertiary structure prediction per-residue confidence estimate (bottom, see methods) suggests a medium confidence total fold (μ-pLDDT: 83.8 ± 13.4), and a high confidence N-terminal “globule” (μ-pLDDT: 90.1 ± 8.7) that is consistently predicted over the top five models (green lines). *domain-A* is consistently predicted without a confident tertiary structure. **b**) Obelisk Oblin-2 has a higher mean normalised conservation (top, 0.26 ± 0.43), and is confidently predicted to form an alpha helix (μ-pLDDT: 97.1 ± 4.6). The Oblin-2 sequence logo (callout, bottom) shows leucine zipper features with “i+7” leucine spacing emphasised in red, with hydrophobic “d” position residues emphasised in yellow (Obelisk-ɑ Oblin-2 sequence shown for reference). Obelisk-ɑ alpha helices (orange boxes, “H” labels), and beta sheets (blue boxes, “S” labels) illustrated for clarity.

**Supplementary Figure 5.**
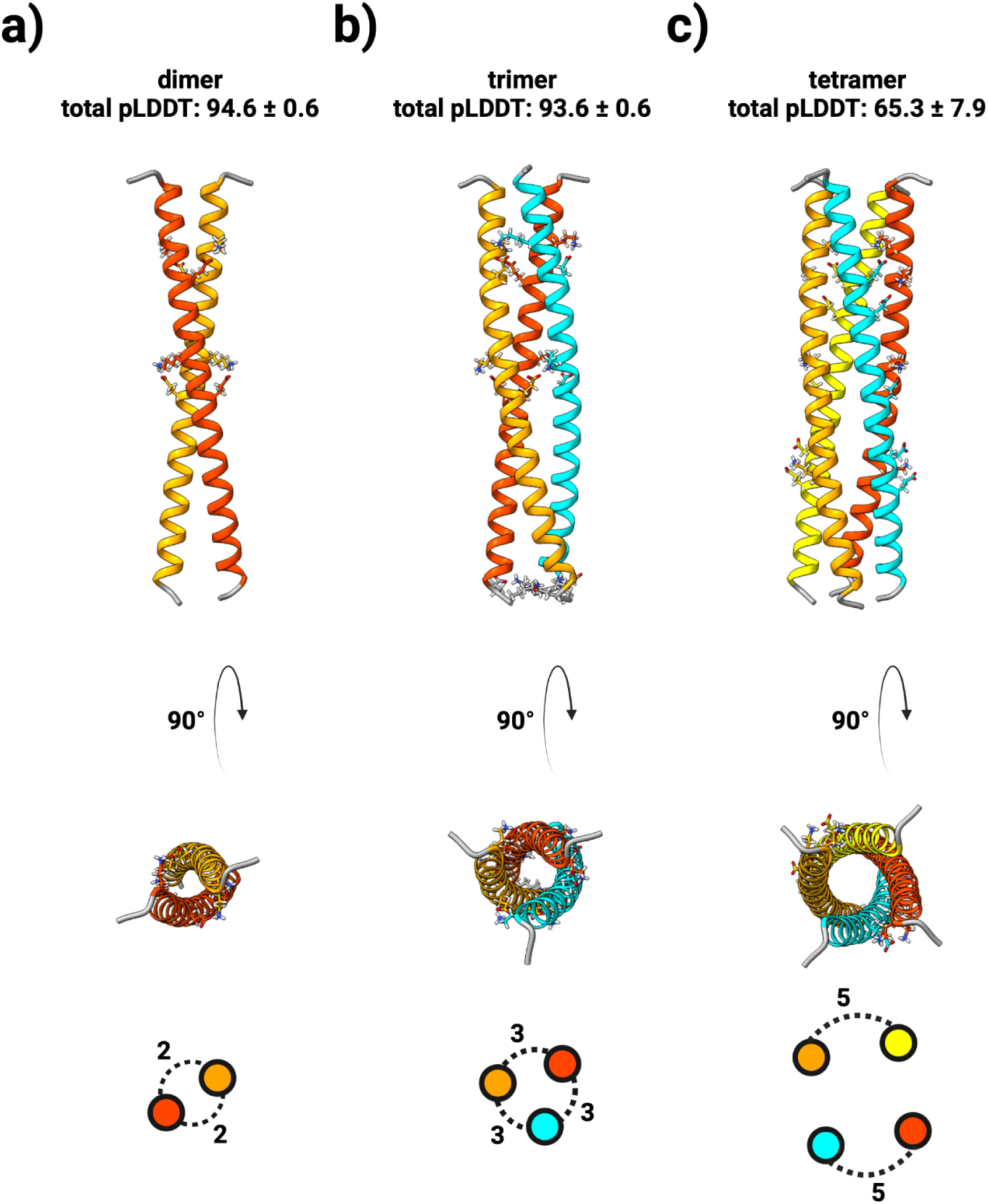
Oblin-2 is predicted to homo-multimerize. tertiary structure predictions of Obelisk-*alpha* open reading frame 2 (Oblin-2) homo-multimers: **a**) dimer (mean pLDDT ± standard deviation: 94.6 ± 0.6), **b**) trimer (mean pLDDT: 93.6 ± 0.6), and **c**) tetramer (mean pLDDT: 65.3 ± 7.9). Residues involved in inter-helix salt bridges emphasised, and salt bridge counts illustrated on bottom.

**Supplementary Figure 6.**
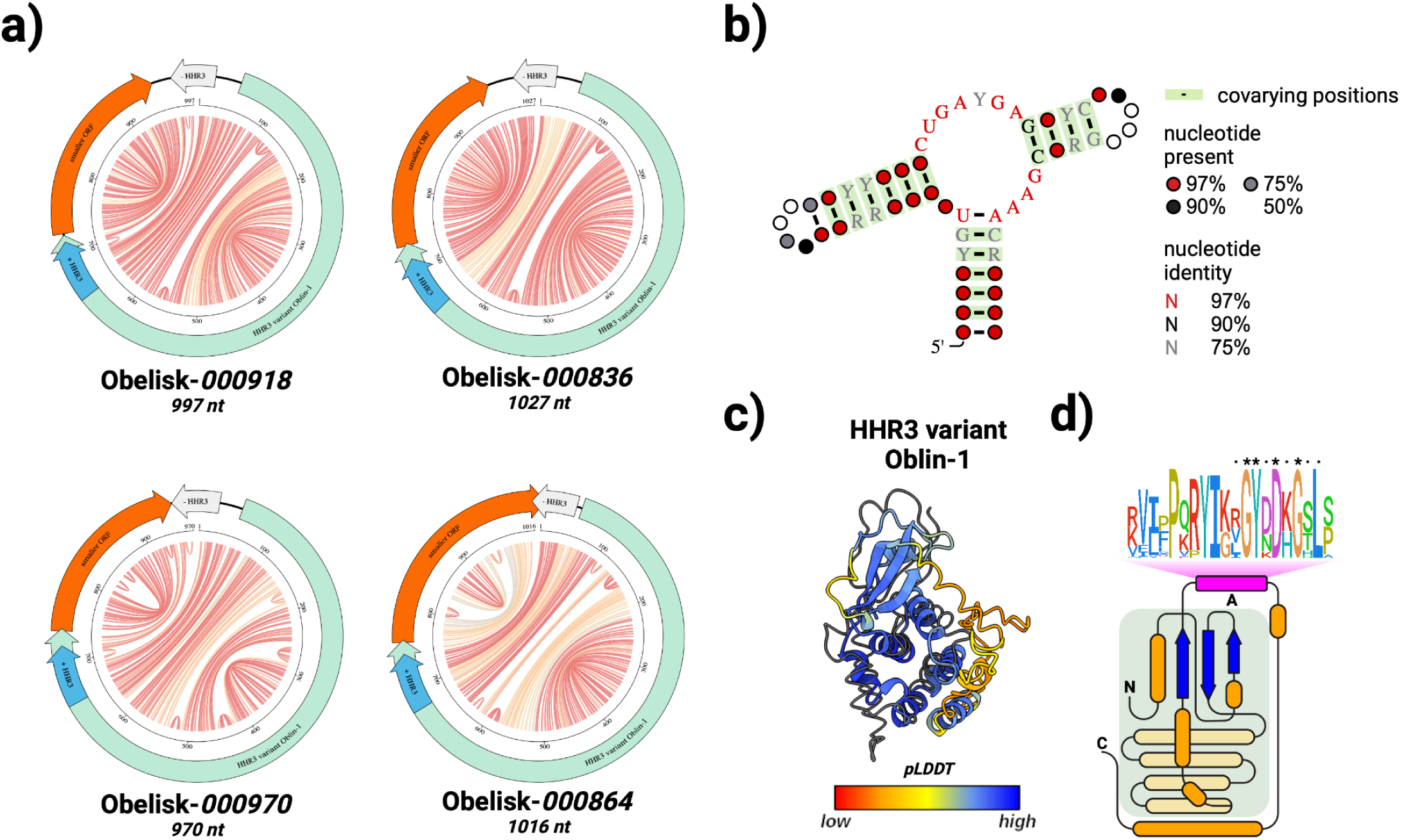
Ribozyme-baring Obelisks encode a diverged Oblin-1. **a**) four “Obelisk-variant hammerhead type-III” (ObV-HHR3) -positive Obelisk genomes from Supplementary Table 1, illustrated as “jupiter” plots where chords represent predicted basepairs (coloured by basepair probability from 0, grey, to 1, red), Oblin-1 homologues illustrated in green, smaller, non-Oblin-2 ORFs in orange, and sense ObV-HHR3 in blue (with antisense ObV-HHR3 in grey). Note the conspicuous placement of ObV-HHR3 relative to Oblin-1 and the smaller ORF. **b**) the `RDVA`-derived, stringently-thresholded ObV-HHR3 covariance model summarised as a secondary structure with bairspair-forming, significantly covarying positions indicated with a green highlight. `IUPAC` “ambiguity codes” ^117^ used to represent RNA diversity: Y = U or C, R = A or G. **c**) `ColabFold` prediction of the “HHR-variant” Oblin-1 tertiary (“*Obelisk_000918*” as the reference sequence) structure built with a custom multiple sequence alignment (MSA) construction (coloured cartoons) superimposed over the `RDVA`-derived MSA prediction for Obelisk-α where possible (black line, Figure 2a, see methods). Prediction confidence (pLDDT) shown as cartoon colouring as in Supplementary Figure 3. **d**) a to-scale (secondary structure) topological representation of “HHR-variant” Oblin-1 with the “globule” shaded in grey (as in Figure 2b), and the *domain-A* emphasised with this bit-score sequence logo (see methods). Conserved “`GYxDxG`” motif emphasised.

**Supplementary Figure 7.**
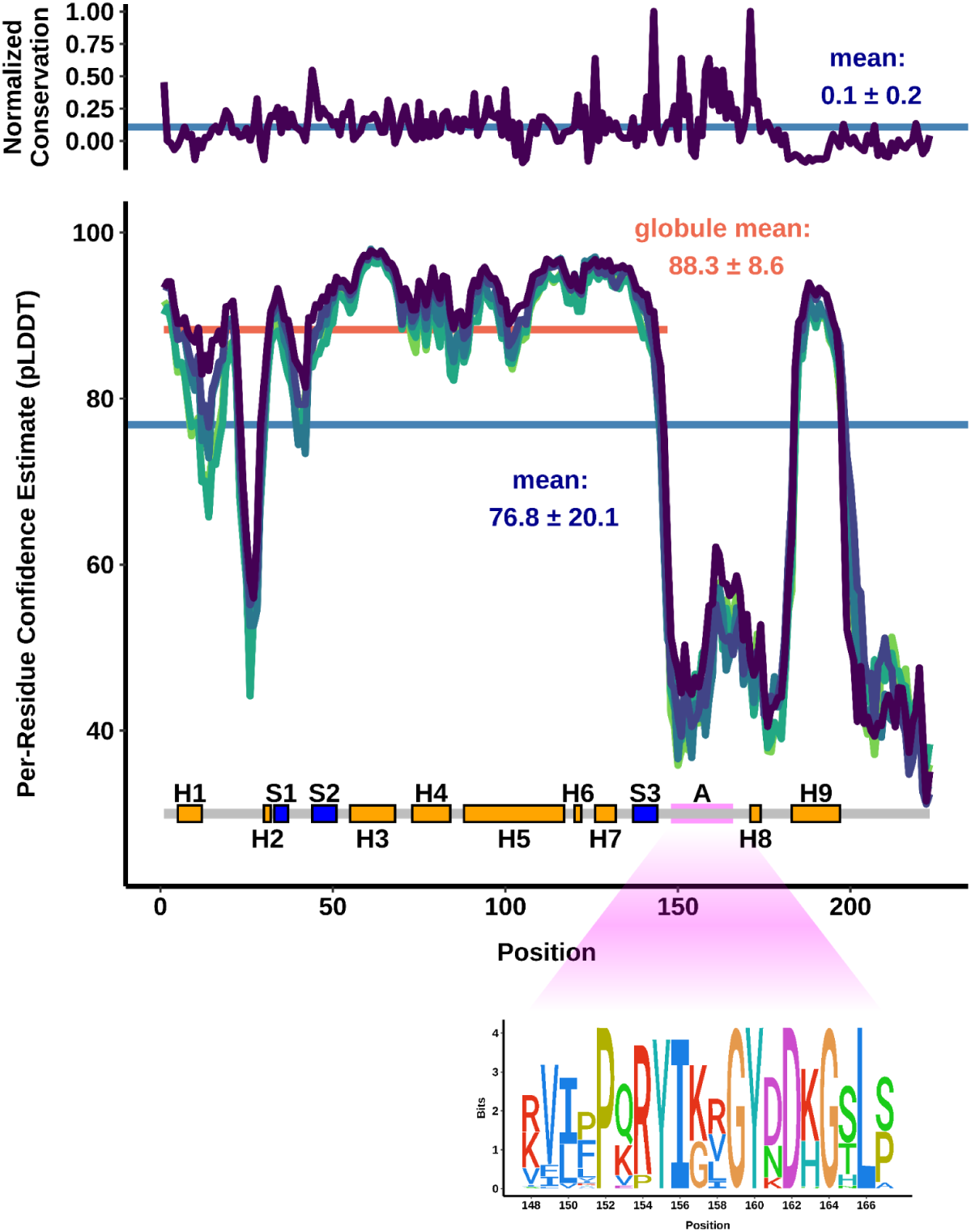
Ribozyme-variant Oblin-1 has similar tertiary fold prediction characteristics to conventional Oblin-1s. normalised conservation (top, above zero = more conserved, see methods) of “Obelisk-variant hammerhead type-III” (ObV-HHR3) “HHR3-variant” Oblin-1 indicates that, similarly to the non-HHR3 Oblin-1 (Supplementary Figure 4), the “HHR3-variant” Oblin-1 is largely poorly conserved (mean normalised conservation ± standard deviation: 0.1 ± 0.2) but retains a conserved *domain-A* (see sequence logo callout, bottom). “HHR3-variant” Oblin-1 tertiary structure prediction per-residue confidence estimate (bottom, see methods) suggests a medium confidence total fold (mean per-residue confidence estimate, μ-pLDDT ± standard deviation of 76.8 ± 20.1), and a higher confidence N-terminal “globule” (μ-pLDDT: 88.3 ± 8.6) that is consistently predicted over the top five models (green lines). *domain-A* is consistently predicted without a confident tertiary structure.

**Supplementary Figure 8.**
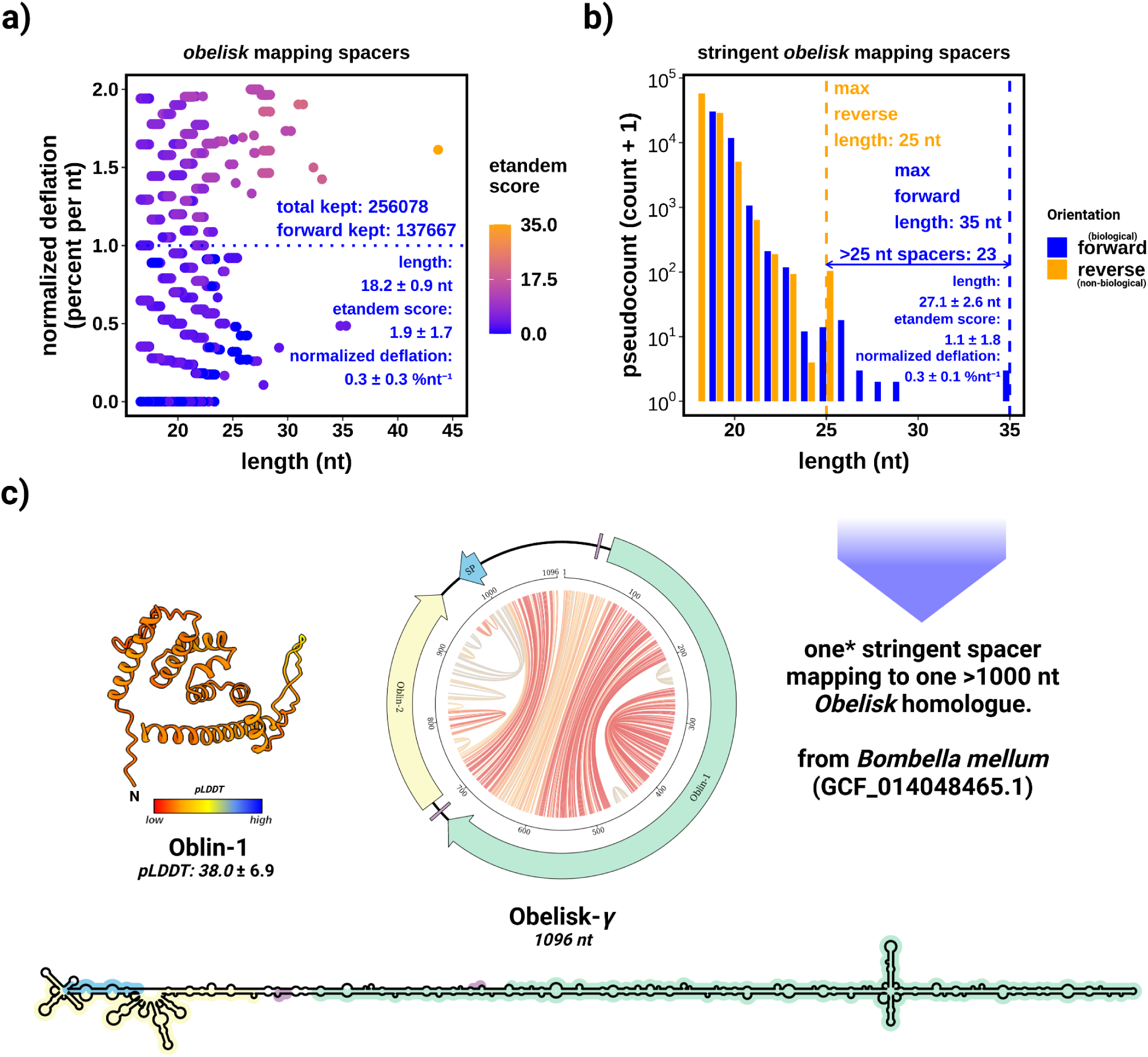
No evidence for capture of Obelisk sequences in available CRISPR-array data. **a**) an x-axis “jittered” scatter plot of Obelisk k-mers that map to the `IMG/M` spacer database ^36^ arranged by a proxy of information content (length-normalised percent deflation, lower = less deflated = more information), coloured by a metric of internal k-mer repetitiveness (see methods). Mappings with a length normalized deflation less than 1.0 percent per nucleotide were kept. Both mappings to “forward” and “reversed” (*not* reverse complemented) Obelisks were kept. Summary statistics on kept k-mers shown in bottom right hand corner. **b**) bar chart representing the noise floor to k-mers kept from a). 23 “forward” mapping k-mers (blue) longer than the longest “reverse” mapping k-mers (orange, 25 nt) were kept. Mappings below this threshold cannot be distinguished from noise. Summary statistics for these kept “forward” k-mers shown in the bottom right hand corner. **c**) ultimately one >1000 nt Obelisk genome was retrieved with two k-mer mappings to the same spacer locus (so the same spacer, see methods). This 1096 nt Obelisk-“gamma” (Obelisk-ɣ) exhibits a “rod-like” predicted secondary structure (“jupiter” plot, centre, “skeleton” diagram, bottom) and contains homologues to Oblin-1 (green) and Oblin-2 (yellow), with the spacer mapping to position ∼1000 (steel-blue “SP” on the jupiter plot). The Obelisk-ɣ Oblin-1 is not predicted to fold into the characteristic “globule” tertiary structure (Figure 4 - tertiary structures). The “frayed” end where the spacer maps deviates from the “rod-ness” of other Obelisks (Figure 4 - “jupiter” plots), suggesting that this Obelisk-ɣ genome might be a chimeric mis-assembly.

**Supplementary Figure 9.**
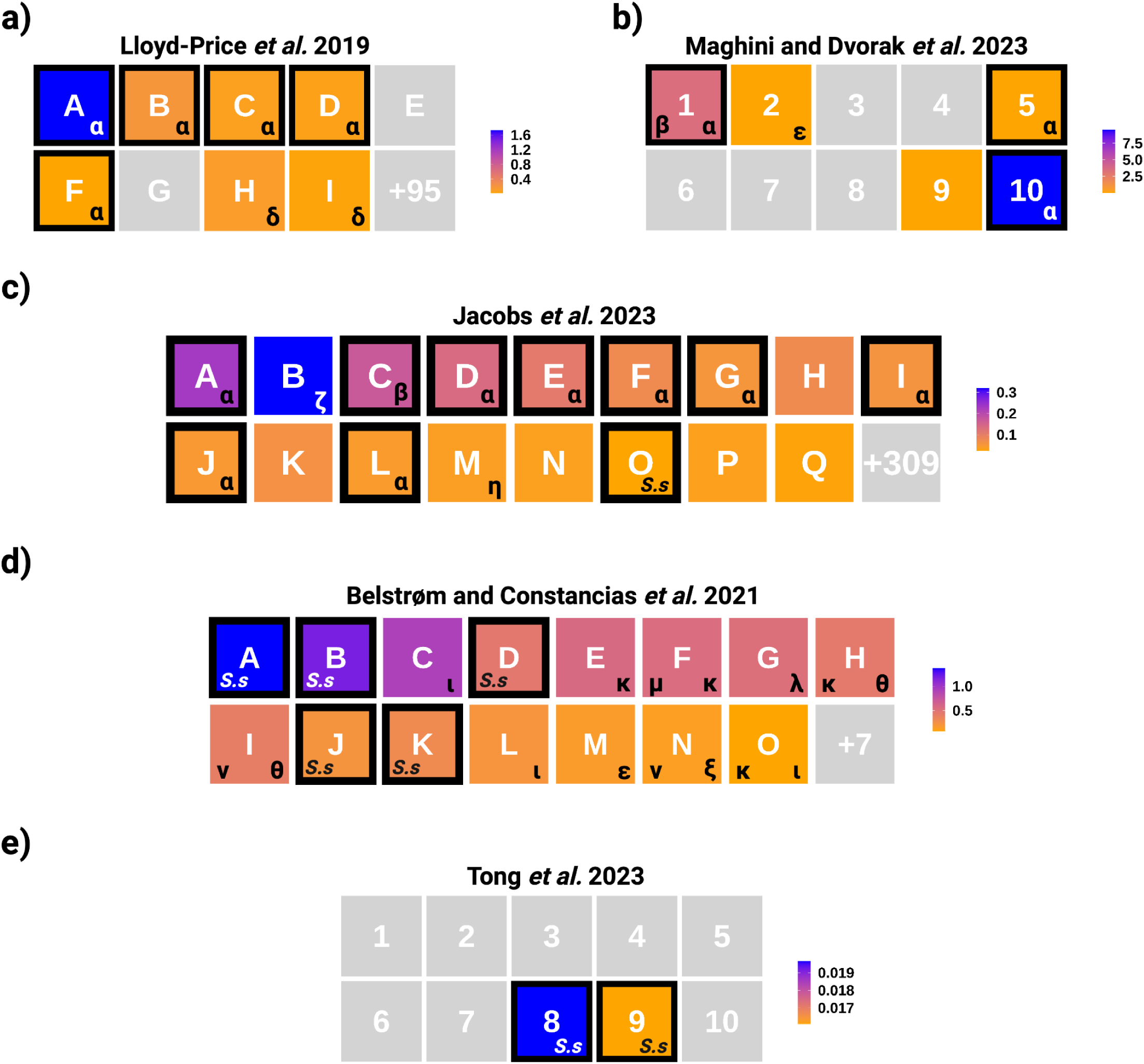
Human gut and oral microbiomes harbour diverse Obelisks. Heatmaps of Obelisk positive donors (>10 reads, averaged over donor if multiple samples) as inferred by k-mer and Oblin-1 pHMM matching (see methods and Table 5, donors with complex internal nomenclature were re-named for clarity see Table 6). Samples emphasised with black boxes were k-mer positive (but not exclusively). Lowercase Greek lettering indicate which Obelisks were found in a given donor as inferred by either k-mer counting (black boxes - k-mer profiling Obelisks -ɑ, -β, and -*S.s*), or by *post hoc* classification of newly assembled and independently clustering Obelisks (see methods). Human gut microbiome samples: **a**) *Lloyd-Price et al. 2019* ^20^, **b**) *Maghini and Dvorak et al. 2023* ^79^, and **c**) *Jacobs et al. 2023* ^112^. Human oral microbiome samples: **d**) *Belstrøm and Constancias et al. 2021* ^37^, and **e**) *Tong et al. 2023* ^113^. Colour scales indicate Obelisk read counts relative to total donor reads ×10^-4^. Greek letter key: α : alpha, β : beta, δ : delta, ε : epsilon, ζ : zeta, η : eta, θ : theta, ι : iota, κ : kappa, λ : lambda, μ : mu, ν : nu, and ξ : xi. Obelisks diagrammed in Figure 4.

**Supplementary Table 1. see Data Availability**

A unified set of Obelisk RNAs grouped hierarchically by percent identity (`circUCLUST` default settings). To ensure stringency, only full length genomes from the `RDVA` dataset were used (subset at 700 nt ≤ length ≤ 2000 nt), as identified by `CircleFinder` (`VNom` settings). Genomes were clustered first at the 80 % identity level, which we define as the boundary between Greek lettering, then at the 95 % identity level, which we define as the sub-type threshold. Open reading frames were then predicted (`prodigal`, `-p meta`) and genomes were converted to match the strand polarity of the largest predicted ORF, placing the first nucleotide of the start codon at the 51st nucleotide. 1,744 80 % identity stringent clusters (composed of 7,202 genomes total) were found. A naming convention is proposed with the following pattern *“Obelisk_X_Y_Z”* where “X” refers to the 80 % cluster ordinate, “Y” to the 95 % cluster ordinate, and “Z” as a unique identifier within the 95 % cluster. The first 15 80 % ordinates are defined as the Obelisks depicted in Figure 4, the next 10 80 % ordinates are defined as the remaining letters in the Greek alphabet (*omicron* through *omega*). As such, the centroid Obelisk-ɑ sequence that is also the centroid of the first 95 % sub-type is defined as “*Obelisk_000001_000001_000001”*. For completeness, an equivalent, additional clustering (see Data Availability) of the `RDVA` dataset without the `CircleFinder`, or `prodigal` steps (subset at 700 nt ≤ length ≤ 1500 nt) is provided. This clustering yielded 6108 80 % clusters of 14,235 genomes total. We caution that this dataset is more likely to be mis-clustered due to unaccounted-for peculiarities of *de novo* assembly, and issues arising from clustering arbitrary reverse-complemented sequences, as such, please use the clusterings (and numberings) in Supplementary Table 1 as the starting point for further Obelisk characterization.

**Supplementary Table 2. see Data Availability**

The *domain-A* alignment and metadata used to construct, and annotate Figure 3.

